# Efficient base editing and development in human embryos without chromosomal alterations

**DOI:** 10.64898/2026.05.30.728989

**Authors:** Stepan Jerabek, Jimin Kim, Julie Sung, Chanju Jung, Marcos Iuri Roos Kulmann, Melisa Isado, Hong-Su Jang, Meng Li, Sakshi Bhatele, Michelle Kappy, Shuangyi Xu, Gue-Ho Hwang, Jia Xu, Diego Marin, Jae-Sun Woo, Sangsu Bae, Nathan Treff, Dieter Egli

## Abstract

Cas9-based tools enable the introduction of genetic lesions to investigate DNA repair outcomes and edit the genome at disease-relevant loci. DNA double-strand breaks (DSBs) induced by CRISPR/Cas9 result in frequent aneuploidy and large deletions, revealing a repair deficiency in early human embryos and limiting the clinical application of this technology. Here we evaluated the DNA repair outcomes of DNA nicks and mismatches introduced using base editors in human embryos at two targets, *PCSK9* and *HBG*. Editing was efficient and, unlike Cas9-induced DSBs, did not result in either chromosomal abnormalities or large deletions. Small insertions or deletions after base editing were rare, and off-target activity was dependent on the guide RNA. Delivering the base editor as a protein at fertilization or at the pronuclear stage allowed normal development to the blastocyst stage and the derivation of edited stem cell lines. In stark contrast, introduction of the editor as RNA resulted in early embryo arrest. Our results demonstrated that, unlike DSBs, DNA nicks and mismatches are efficiently repaired in human embryos, allowing specific on-target changes without genotoxic consequences.

## INTRODUCTION

Gene editing systems rely on cellular repair machinery to correct DNA breaks, nicks, mismatches, or abasic sites. This repair may be cell type- and species-specific and is thus important to understand in different contexts. Previous studies using CRISPR/Cas9 in human embryos uncovered the deficiency of DNA DSB repair, resulting in aneuploidies consisting of segmental as well as whole chromosome errors in up to half of all cells^1,2^. Chromosome breaks, genome instability, and aneuploidy occur spontaneously in human embryos and are thought to be a common cause of developmental failure and genetic disease but are still poorly understood. Therefore, the experimental induction of defined and targeted genetic lesions is a powerful way to investigate DNA repair outcomes and to understand the consequences of DNA damage in the earliest stages of human development.

In addition to chromosome loss, additional on- and off-target genotoxic consequences of Cas9-induced DNA DSBs were reported in mouse and human cells, including large deletions, chromosome truncation, complex chromosomal rearrangements, and chromothripsis^3–9^. Large deletions after CRISPR/Cas9 targeting have also been detected in mouse and human embryos^10–12^. Regarding potential human heritable gene editing, the various detrimental outcomes of DNA DSBs in human embryos, which are also challenging to accurately characterize, make utilization of CRISPR/Cas9 a highly implausible proposition.

Alternatively, base editors use Cas9 nickase (nCas9) to generate only a single-strand break (SSB) in the target DNA locus, thus enabling the study of DNA repair outcomes of targeted nicks and mismatches. Although a DSB is not mandatory, when a DNA replication fork encounters a SSB, it can result in a DNA DSB^13^. Through this conversion, nCas9 also carries the potential for induced insertions and deletions (indels), large deletions, and chromosomal changes. Editing by both adenine base editors (ABEs) and cytidine base editors (CBEs) can lead to indels with frequencies depending on the specific variant^14–17^. DNA DSB can also be created by error-prone repair of an abasic site or by bystander editing in the target window^17,18^. Recent studies focusing on the genotoxicity of genome editing tools in human cells have found occasional large deletions (> 100 bp) after editing by both ABEs and CBEs^19^ and translocations after multiplex CBE (but not ABE) editing of different chromosomes^20^. Altogether, these studies establish that base editors introduce potentially dangerous DNA lesions. As such lesions also occur spontaneously, the knowledge gained through studying the genetic and genomic consequences of base editing in early human embryos is relevant to both normal development and to gene editing.

Here we evaluated the repair outcomes of DNA nicks introduced by Cas9 nickase and mismatches associated with ABEs at two different genomic sites, the proprotein convertase subtilisin/kexin type 9 (*PCSK9)* gene and gamma globin genes (*HBG1*/*2*), in early human embryos. We chose these genes because they are well studied in the context of somatic gene editing, rather than based on therapeutic promise in the germ line. We established that base editing at the one-cell stage is efficient and precise even when two genes are targeted simultaneously. Importantly, no segmental aneuploidies or large deletions were observed after base editing but were frequent after Cas9-induced DSBs at the same genomic site. This enabled the derivation of high-quality blastocysts and human embryonic stem cells (hESCs) deficient in *PCSK9* and devoid of detectable genotoxic consequences. Interestingly, normal development only occurred when the base editor was introduced as a ribonucleoprotein (RNP). Injection of the in vitro transcribed mRNA led to developmental arrest between the 1-4 cell stage, pointing to a hitherto unknown mechanism of RNA-mediated embryo arrest in human.

## RESULTS

### Efficient and precise editing of *PCSK9* and *HBG* genes in human embryos

To investigate DNA repair of SSBs in human embryos, we selected two sgRNAs that target *PCSK9* or *HBG1/2* located on human chromosomes 1 and 11, respectively **(Fig. 1a, Extended Data Table 1)**. Both sgRNAs have been previously validated in somatic cells^15,21^. The *PCSK9* sgRNA was designed for ABE-mediated A→G edit at the splice site between exon 1 and intron 1 of the *PCSK9* gene, resulting in read-through and premature termination at the TAG stop codon 10 bp from the end of exon 1^21^. This mutation reproduces the effect of naturally occurring nonsense mutations in *PCSK9* leading to low-density lipoprotein cholesterol and reduced risk of coronary heart disease^22^, but is not known to occur naturally. The *HBG1/2* sgRNA was designed to enable the A→G substitution at the -198 position in the promoter region of the *HBG1* and *HBG2* genes^15^. This naturally existing variant, known as the British mutation, leads to fetal hemoglobin expression in adulthood, ameliorating the symptoms of β-thalassemia and sickle cell disease^23^. Both sgRNAs delivered with ABE mRNA in human embryonic kidney 293 (HEK293T) cells or hESCs showed high editing efficiency between 70% to over 95% **(Extended Data Fig. 1a-d)**.

**Figure 1.**
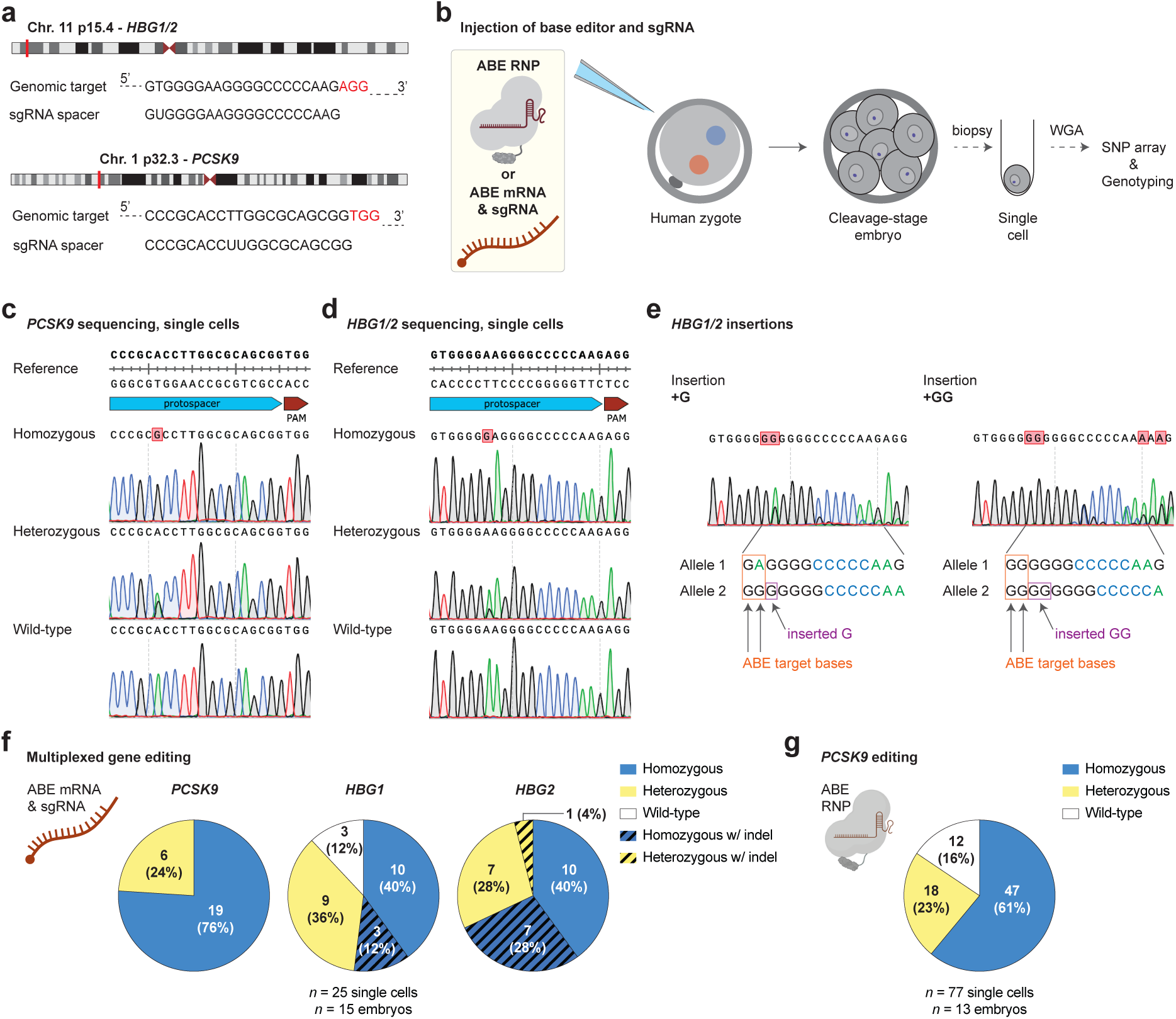
Efficient and precise gene editing by ABE after injection in human zygotes. **a,** Schematic illustrating genomic location of two target genes, *PCSK9* and *HBG1/2*, and sequences of used sgRNAs. **b,** Workflow for ABE mRNA or ABE ribonucleoprotein (RNP) delivery into two-pronuclear (2PN) human zygotes and analysis of whole-genome-amplified (WGA) samples from single cells collected from injected embryos. **c-e,** Sanger sequencing chromatograms showing base editing outcomes at two targeted loci, *PCSK9* (**c**) and *HBG1/2* (**d, e**). Two types of insertions detected at *HBG1/2* are depicted (**e**) with Sanger sequencing chromatograms and nucleotide base sequences according to the analysis using EditR^48^. **f, g,** Quantification of editing outcomes based on Sanger sequencing of single cells isolated from ABE mRNA- (**f**) and RNP-edited (**g**) human embryos.

We then continued to analyze ABE editing in human embryos **(Fig. 1b)**. To maximize the information to be gained, we injected ABE mRNA along with both *PCSK9* and *HBG1/2* sgRNAs into human two-pronuclear (2PN) zygotes and performed Sanger sequencing of individual blastomeres **(Fig. 1c-e, Supplementary Table 1)**. Of 25 blastomeres from 15 embryos, cells homozygous for the intended edit accounted for 76% (19/25) at *PCSK9*, 52% (13/25) at *HBG1*, and 68% (17/25) at *HBG2* **(Fig. 1f)**. Of 11 indels found in *HBG1/2*-edited samples, eight contained the insertion of a single guanine, and three showed insertions of two additional guanines **(Fig. 1e)**. Neither bystander edits nor indels were detected around the *PCSK9* target site. This corresponds with our results from experiments in cultured cells, in which *PCSK9* sgRNA was the better of two used guides in terms of efficiency (∼96%) and indel frequency (less than 0.5%) **(Extended Data Fig. 1a-d)**, and we detected rare, short on-target indels only after editing with *HBG1/2* sgRNA **(Extended Data Fig. 1e, f)**.

For additional human embryo experiments, we purified the ABEmax protein **(Extended Data Fig. 1g)**, allowing us to compare the ABE RNP delivery with mRNA injections, for which the number of blastomeres per embryo developed within the first three days was very small **(Fig. 1f)**. Sanger sequencing analysis of 77 blastomeres from 13 human embryos identified 61% (47/77) cells with homozygous edits, 23% (18/77) cells with a heterozygous edit, and 16% (12/77) wild-type cells **(Fig. 1g)**. Similar to the previous mRNA-based editing, no bystanders or indels were found at *PCSK9* loci in any of the sequenced blastomeres. In conclusion, ABE injections into 2PN human zygotes as mRNA or RNP enable efficient and precise target gene editing, with ABE RNP-injected embryos showing better early development.

### Base editing does not lead to large deletions at on-target sites in human embryos

To compare CRISPR/Cas9 to base editing in early human embryos, we collected 24 cells from seven cleavage-stage embryos injected with Cas9 RNPs consisting of both *PCSK9* and *HBG1/2* sgRNAs. Amplicon next-generation sequencing (NGS) data analysis of pooled whole-genome amplified (WGA) samples showed that targeting efficiencies reached over 70% for *PCSK9* and about 45% for *HBG1/2,* with Cas9-induced deletions of various sizes **(Extended Data Fig. 2a)**. For NGS analysis of base-edited cells, we pooled samples previously analyzed by Sanger sequencing **(Fig. 1f, g)** with additional WGAs. We combined a total of 124 (eight multicell) samples from 40 embryos for evaluation of *PCSK9* editing and 27 (one multicell) samples from 17 embryos for evaluation of *HBG1/2* editing **(Supplementary Table 1)**. Our analysis showed high base-editing efficiencies, 65% and 52% for *PCSK9* and *HBG1/2*, respectively **(Extended Data Fig. 2b)**. However, deletions were far more frequent in CRISPR/Cas9-treated samples when compared to base-edited samples at both *PCSK9* and *HBG1/2* loci (*p* < 0.00001 for both target loci; Fisher’s exact test).

CRISPR/Cas9 can induce large deletions in mouse and human embryos^10,12^. We used long-range PCRs **(Fig. 2a, b, Extended Data Fig. 2c)** to screen for large deletions in 24 single-cell samples from seven CRISPR/Cas9-injected human embryos. Two samples produced a single, shorter PCR band for *PCSK9* and *HBG2* **(Fig. 2c)**, with deleted regions of 740 bp and 1116 bp, respectively, instead of the expected ∼1.8-kilobase (kb) PCR amplicon **(Fig. 2d)**. More than one-third of all amplifications (17/48; 35%) from CRISPR/Cas9-edited embryos lacked a PCR product, which may be due to: i) chromosomal change resulting in the loss of the targeted locus, ii) deletions even longer than the ∼1.8-kb analyzed region, leading to removal of primer binding sites; or iii) the presence of an unrepaired DNA DSB or other rearrangements not detectable by PCR.

**Figure 2.**
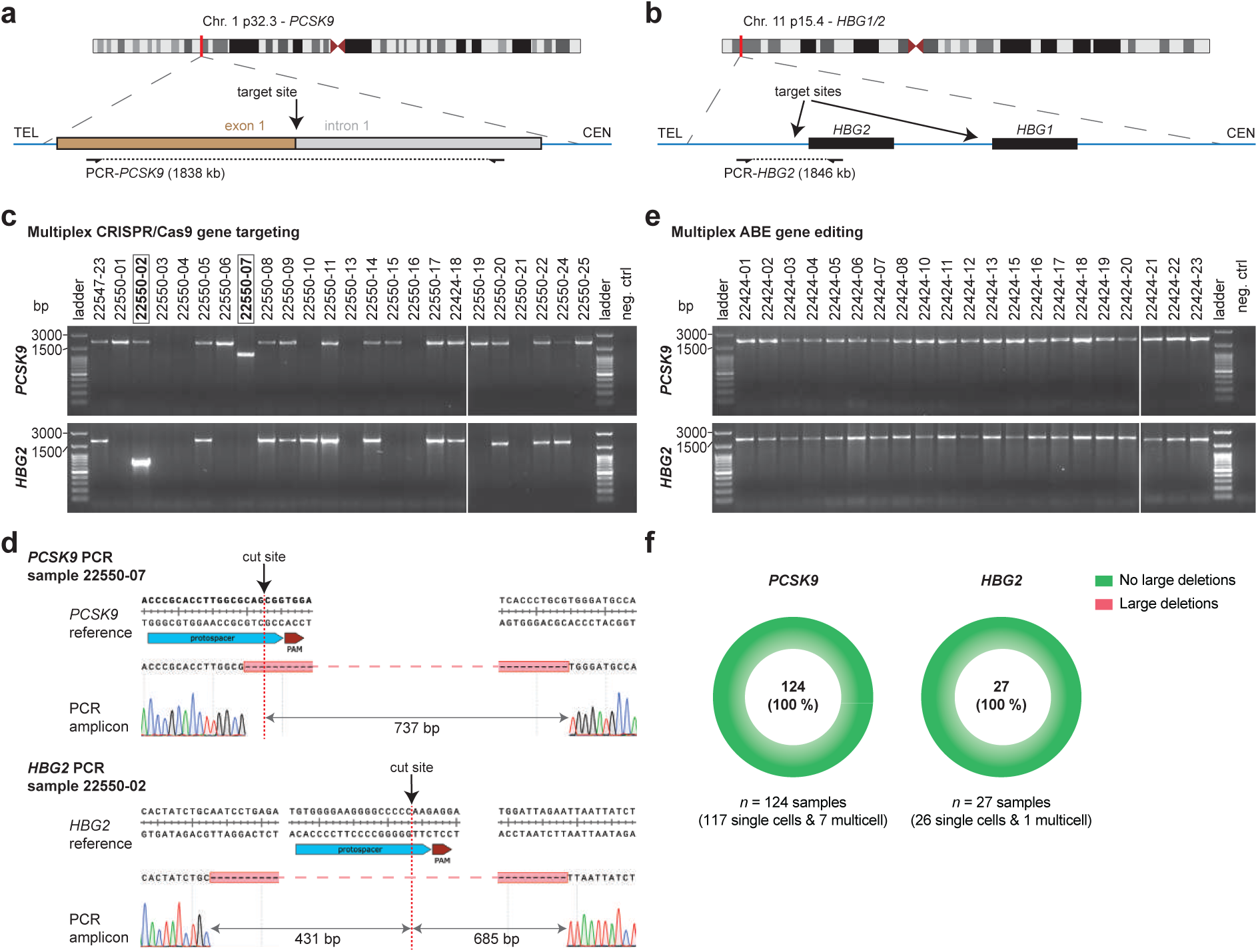
Analysis of large deletions at target sites in human embryos after zygote injections of CRISPR/Cas9 or adenine base editor. **a, b,** Design of PCRs for two genes, *PCSK9* (**a**) and *HBG2* (**b**), targeted by CRISPR/Cas9 and ABE. Primers were designed to amplify ∼1.8-kb-long genomic regions with the target sites being approximately in the middle of each PCR amplicon. **c,** Agarose gel electrophoretic analysis after PCR amplifications of *PCSK9* (upper panel) and *HBG2* (lower panel) target sites using WGA blastomere samples from human embryos injected at the zygote stage with CRISPR/Cas9 RNPs including both *PCSK9* and *HBG1/2* sgRNAs (multiplex gene targeting). Black boxes highlight two samples for which shorter-then-expected fragments were detected, indicating the presence of large deletions. **d,** Sanger sequencing-based verification of Cas9-induced large deletions at *PCSK9* (upper panel) and *HBG2* (lower panel) target sites in two samples highlighted in panel (**c**). **e,** Agarose gel electrophoretic analysis after PCR amplifications of *PCSK9* (upper panel) and *HBG2* (lower panel) target sites using WGA blastomere samples from human embryos injected at the zygote stage with ABE mRNA and both *PCSK9* and *HBG1/2* sgRNAs (multiplex gene editing). **f,** Quantification based on PCR analyses of target sites in samples from ABE-injected human embryos. No large deletions were identified for *PCSK9* or *HBG2* loci.

All PCR reactions for 124 samples from *PCSK9* base-edited embryos and 27 samples from *HBG1/2* base-edited embryos showed normal-sized amplicons **(Fig. 2e, f)**. The differences between CRISPR/Cas9-targeted and base-edited samples in amplification efficiency and in the frequency of large deletions were both significant (*p* < 0.00001 and *p* = 0.028, respectively; Fisher’s exact test). In conclusion, adenine base editing shows far more specificity and reduced complexity of genetic modifications in preimplantation human embryos when compared to DNA DSB repair outcomes after CRISPR/Cas9 targeting.

### Chromosomal integrity after adenine base editing in human embryos

Cas9-induced DNA DSBs in human embryos cause frequent segmental or whole chromosome changes^1,2,11^. We used single nucleotide polymorphism (SNP) arrays and copy number analysis to detect aneuploidies in single cells from CRISPR/Cas9- or ABE-injected human embryos to directly compare chromosomal consequences of either technology at the same genomic sites **(Fig. 3a)**.

**Figure 3.**
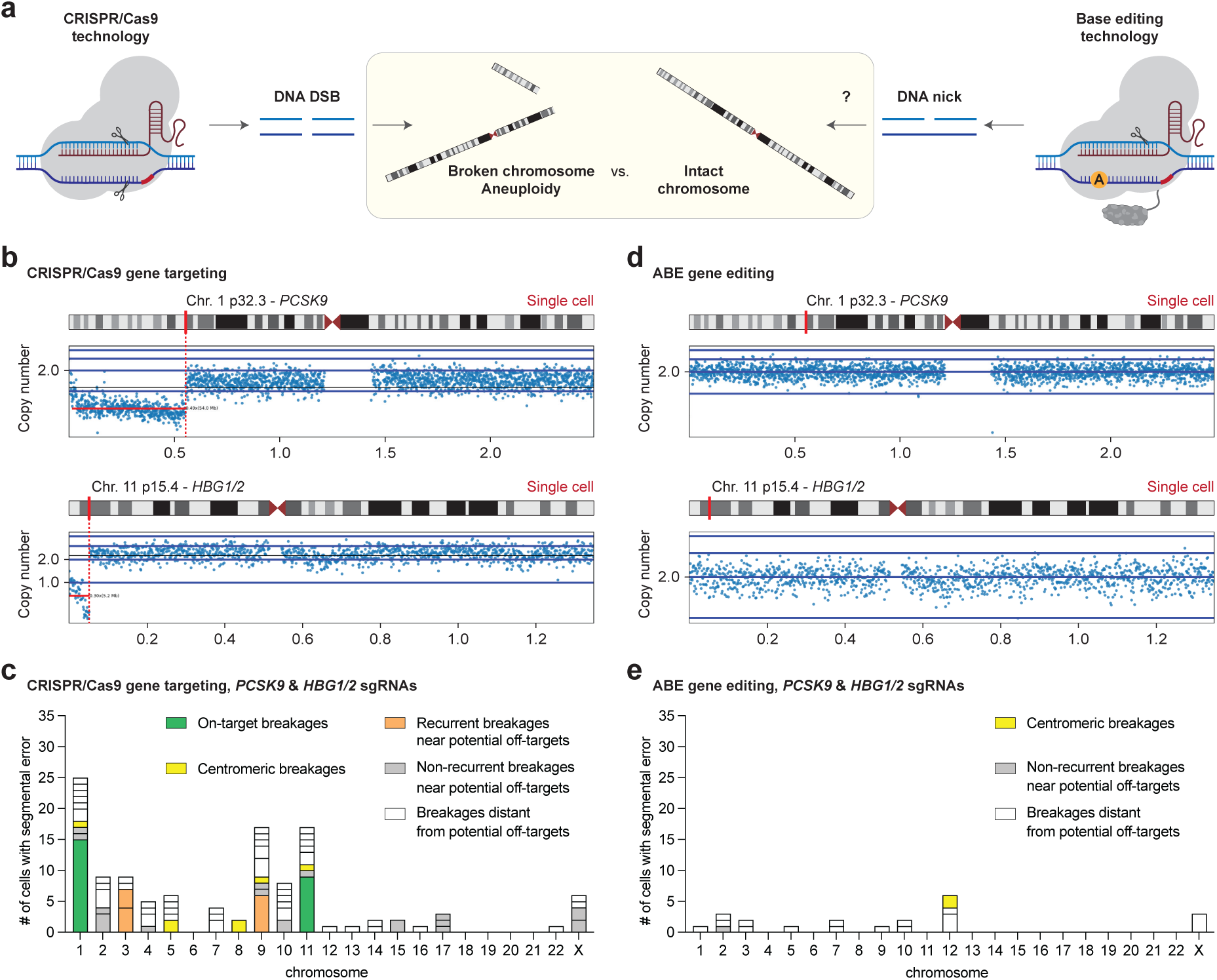
Base editing in early human embryos without chromosome changes. **a,** Schematic illustrating the consequences of Cas9-generated DNA DSBs on genome stability and the hypothesis about the absence of chromosome breaks after adenine base editing involving DNA nicks in human embryos. **b,** Copy number plots for chromosomes 1 (upper panel) and 11 (lower panel) in single blastomeres from CRISPR/Cas9-injected human embryos. Positions of target genes, *PCSK9* and *HBG1/2*, on their respective chromosomes are highlighted. **c,** Summary of the chromosome break sites mapping in cells collected from CRISPR/Cas9-injected human embryos. Copy number plots and SNP array data sets were analyzed for *n* = 23 cells from CRISPR/Cas9-injected embryos. Individual breakages are categorized based on their position relative to on-target sites and off-targets predicted by CRISPOR^46^. **d,** Copy number plots for chromosomes 1 (upper panel) and 11 (lower panel) in single blastomeres from ABE-injected human embryos. **e,** Summary of the chromosome break sites mapping in cells collected from ABE-injected human embryos. Copy number plots and SNP array data sets were analyzed for *n* = 117 cells from ABE-injected embryos.

We analyzed 23 WGA blastomere samples from human embryos injected with CRISPR/Cas9 RNPs with *PCSK9* and *HBG1/2* sgRNAs and mapped genomic coordinates of chromosome break sites (**Supplementary Table 2**). Out of 119 breakages, 25 mapped to the targeted loci; 15 were detected at *PCSK9* on chromosome 1, and nine were detected at *HBG1/2* on chromosome 11 **(Fig. 3b, c)**. In addition to on-target Cas9-induced segmental changes, we identified spontaneous break sites, as well as 18 breakages proximal to in silico-predicted off-targets for *HBG1/2* sgRNA **(Fig. 3c, Extended Data Fig. 3a-c, Extended Data Table 2)**. No chromosomal breakages mapped near in silico predicted off-targets for *PCSK9*.

Chromosomal analysis of 117 WGA blastomere samples from *HBG1/2* and/or *PCSK9* base-edited human embryos showed that chromosomes 1 and 11, on which the two target sites are located, remained intact in every single cell **(Fig. 3d, e, Extended Data Fig. 3d, e, Supplementary Table 2)**. Only one of 21 segmental copy number transitions mapped within a 200-kb window from the predicted off-target site on chromosome 2; however, no Cas9-induced breakages were observed for this site. Break sites in ABE-injected human embryos were located predominantly in gene-poor regions, as characteristic for spontaneous chromosome breakages^24^ **(Extended Data Fig. 3f, Supplementary Table 3)**. In conclusion, genome integrity at the target site is preserved in ABE-injected human embryos.

### Normal development after base editing in human zygotes

We asked whether injection of ABE editor in the form of mRNA or a protein would impact the normal development of human embryos. We evaluated two time points of delivery: co-injection of ABE with sperm at fertilization and injection into fertilized, 2PN embryos **(Fig. 4a)**. Of the 19 human embryos injected with ABE mRNA at the 2PN stage, none (0/19; 0%) developed beyond the early cleavage stage **(Fig. 4b)**. In contrast, after injection of ABE RNPs at the 2PN stage, significantly more embryos (8/25; 32%) reached the morula and the blastocyst stages. ABE RNP injections were also compatible with normal development when performed in metaphase II (MII) oocytes at fertilization, as demonstrated by four high-quality blastocysts (4/12; 33%) **(Fig. 4b, Extended Data Table 3)**.

**Figure 4.**
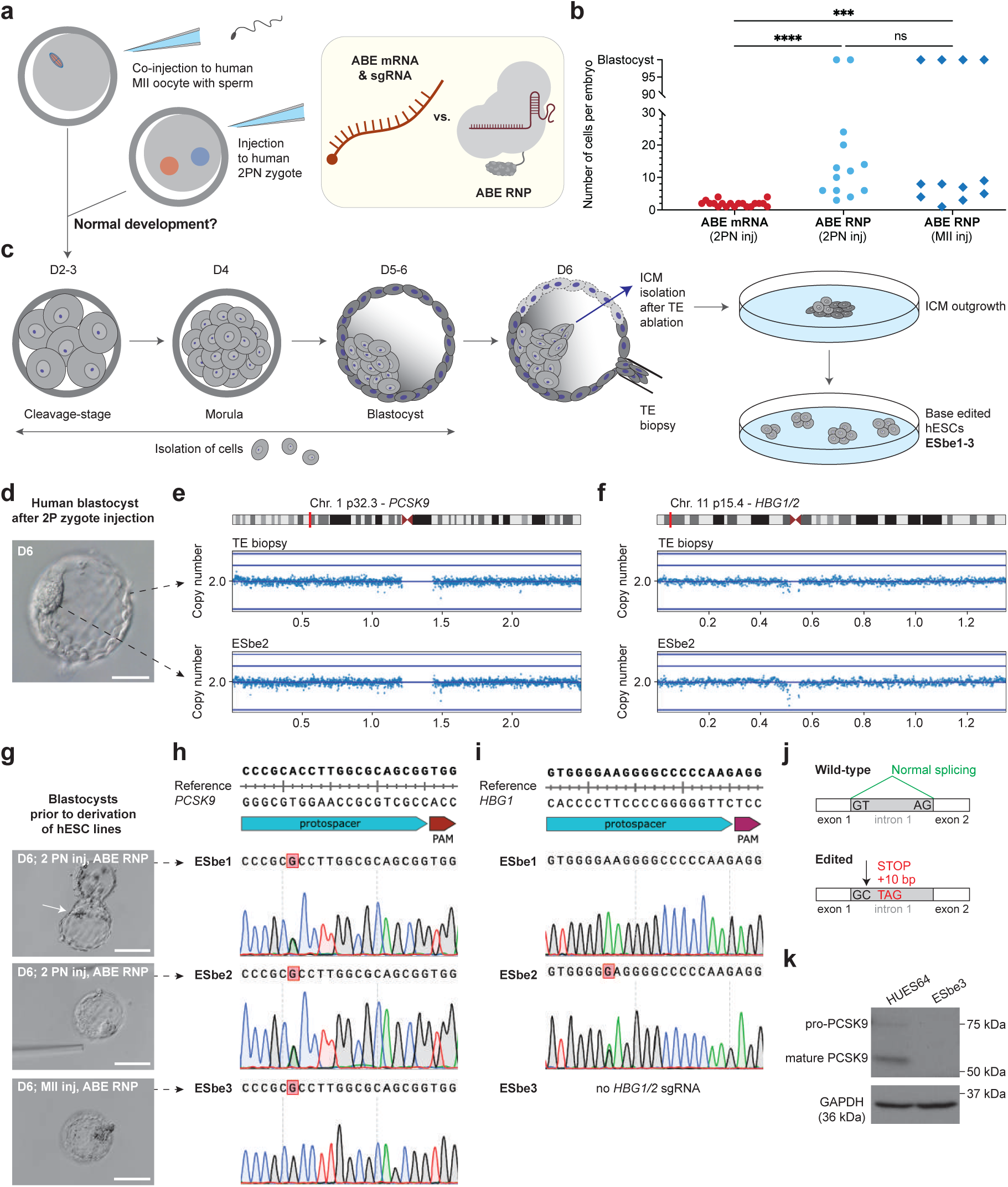
Normal development of base-edited human embryos. **a,** Schematic showing injection of base editors as mRNA or a purified protein into metaphase II (MII) oocytes with a sperm or into two-pronuclear (2PN) zygotes. **b,** Developmental potential of human embryos assessed by the number of cells after injections of adenine base editor (ABE) encoded by mRNA (*n* = 19) or ABE ribonucleoprotein (RNP) (*n* = 25). Within the ABE RNP cohort, *n* = 12 embryos were injected at fertilization (dark blue diamonds). Statistical significance established with one-way ANOVA. **c,** Schematic illustrating preimplantation development of ABE-injected embryos, trophectoderm (TE) biopsy from day 6 (D6) blastocyst, and derivation of human embryonic stem cells (hESCs) from the inner cell mass (ICM). **d,** A blastocyst injected with ABE RNP at the 2PN zygote stage. Scale bar, 100 µm. **e, f,** Copy number plots showing the integrity of targeted chromosomes 1 (**e**) and 11 (**f**) in TE biopsy (upper panel) and derived hESCs (lower panel). **g,** Blastocysts developed after ABE RNP injections into 2PN zygotes or an MII oocyte at fertilization. Scale bars, 200 µm. **h, i,** Sanger sequencing chromatograms showing editing outcomes at *PCSK9* (**h**) and *HBG1* (**i**) in three hESC lines from ABE-injected embryos. Scale bars, 200 µm. **j, k,** Schematic showing the effect of adenine base editing of *PCSK9* exon 1 splice-donor as predicted by Musunuru et al.^21^ (**j**) and a Western blot showing complete PCSK9 knockout in hESCs with homozygous *PCSK9* edit (**k**). GAPDH was used as the loading control.

We derived hESC lines from three blastocysts developed from ABEmax RNP base-edited embryos: two lines (ESbe1 and ESbe2) from embryos injected as 2PN zygotes and one additional hESC line (ESbe3) from an embryo injected at fertilization **(Fig. 4c)**. SNP array data obtained for trophectoderm (TE) biopsies and newly derived hESCs allowed us to track genetic relatedness between identical-embryo samples **(Extended Data Fig. 4a)** and confirm balanced copy number of targeted chromosomes 1 and 11 **(Fig. 4d-f)**. For the *PCSK9* target locus, ABE injections into human embryos resulted in heterozygous editing in ESbe1 and ESbe2 and homozygous editing in ESbe3 **(Fig. 4g, h)**. Sanger sequencing of *HBG1/2* target genes revealed heterozygous editing in ESbe2 **(Fig. 4i)**.

All three base-edited hESC lines derived from ABE RNP-injected embryos have normal karyotype and express key pluripotency markers, SOX2 and OCT4 **(Extended Data Fig. 4b, c)**. A→G editing at the splice donor site after exon 1 leads to premature termination of translation^21^ **(Fig. 4j)**. Indeed, we found that homozygous editing effectively removes PCSK9 protein in ESbe3 hESC line **(Fig. 4k)**.

In injected 2PN zygotes (day 1), chromosomes have already been replicated at the time of ABE delivery, and a total of four *PCSK9* targets are present, increasing the likelihood for genetic mosaicism. We found mosaicism at the *PCSK9* loci in seven out of nine human cleavage-stage embryos (7/9; 78%) from which we collected single blastomeres at the four-cell stage or later; only two embryos (#10 in group BE1 and #4 in group BE3) showed uniform editing **(Extended Data Fig. 4d, Supplementary Table 1)**. We also performed NGS using seven multicell samples that developed beyond the cleavage stage. The TE biopsy from the 5AA embryo, from which the ESbe2 hESC line was derived, was uniformly heterozygous. Two samples, a complete morula and a complete blastocyst-stage embryo, showed uniform homozygous *PCSK9* base editing. We detected impure on-target *PCSK9* editing (2.6% NGS reads) only in one sample from a 20-cell embryo **(Extended Data Fig. 4e, Supplementary Table 4)**. Our current dataset in MII oocytes does not allow conclusive assessment of mosaicism after ABE delivery at intracytoplasmic sperm injection (ICSI). In summary, ABE injections into human 2PN embryos or at fertilization are compatible with normal development to blastocysts and successful derivation of base-edited hESCs.

### Cas9-induced chromosome break sites predict off-target base editing

To assess the potential for off-target editing in human embryos, we analyzed genomic sites for which we observed chromosomal breaks proximal to in silico-predicted off-targets for the *HBG1/2* sgRNA **(Fig. 3c, Extended Data Table 2)**. For this, we pooled 24 and 27 single-cell WGA samples from CRISPR/Cas9- and ABE-injected embryos, respectively. We then performed amplicon NGS of 14 candidate off-target regions (OFF1-14) using pooled CRISPR/Cas9 and ABE samples for side-by-side comparisons of DNA repair outcome at each genomic region **(Extended Data Fig. 5a)**. Cas9-generated indels were found at 11 out of 14 analyzed sites **(Fig. 5a, b, Extended Data Fig. 5b-n, Supplementary Table 5)**, confirming that most potential off-target loci determined through Cas9-induced chromosome break mapping are indeed real off-targets for *HBG1/2* sgRNA. The efficiency of CRISPR/Cas9 targeting at off-target sites (OFF1, OFF5, OFF7, and OFF8) was comparable to the on-target *HBG1/2* site **(Fig. 5b)**, despite up to four mismatches with the sgRNA spacer sequence, exceeding the three-mismatch cutoff commonly applied by studies relying solely on predictive algorithms.

**Figure 5.**
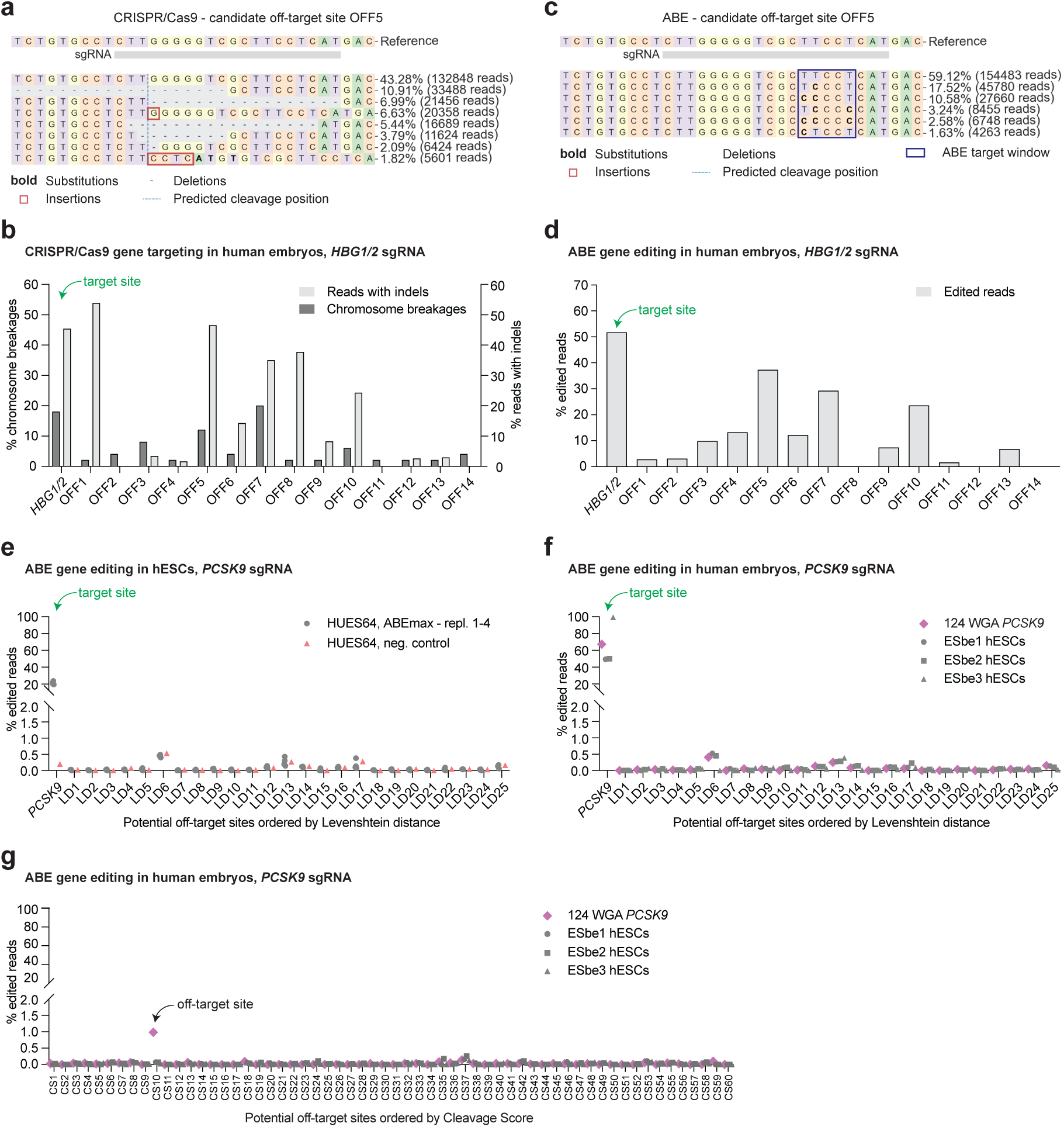
Assessment of off-target editing in human embryos and in human embryonic stem cells derived from base-edited embryos. **a, b,** Alleles frequency tables around a potential off-target site OFF5 after NGS using a pooled WGA sample from CRISPR/Cas9-targeted (**a**) and ABE-edited (**b**) human embryos. For both technologies, multiplexed editing was performed using *PCSK9* and *HBG1/2* sgRNAs. *n* = 24 WGA blastomere samples were pooled for CRISPR/Cas9 NGS analysis, and *n* = 27 WGA blastomere samples were pooled for ABE NGS analysis. Chromosomal breakages at off-target sites were evaluated based on SNP arrays for individual blastomeres. **c, d,** The efficiency of CRISPR/Cas9 targeting (**c**) and ABE editing (**d**) at *HBG1/2* and candidate off-targets based on chromosome breakage analysis. For CRISPR/Cas9 targeting, the percentage of segmental chromosome breaks and the percentage of NGS reads with Cas9-generated indels are shown for each genomic site. **e, f,** ABE editing efficiency at *PCSK9* and potential off-targets (ordered by Levenshtein distance; LD1-25) in transfected HUES64 hESCs (*n* = 4) and untreated control (**e**), and in human embryos and three hESC lines derived from base-edited embryos (**f**). *n* = 124 WGA blastomere samples were pooled for the NGS analysis of individual sites. Signals at and below 0.5% were considered background, as clonal lines and embryo samples produced identical results. **g,** ABE editing efficiency at potential off-targets (ordered by cleavage score; CS1-60) in human embryos and three hESC lines derived from base-edited embryos. *n* = 124 WGA blastomere samples were pooled for the NGS analysis.

NGS analysis of the same sites in ABE-injected embryo samples revealed base editing at 11 out of the same 14 candidate off-targets **(Fig. 5c, d, Extended Data Fig. 6a-m, Supplementary Table 5)**. The most active ABE off-target site (OFF5) was edited in approximately 37% of analyzed reads, only modestly lower than on-target *HBG1/2* editing **(Fig. 5 d)**. These results show that Cas9-induced chromosome breakage analysis in human embryos also enables the identification of active ABE off-target sites. Though off-target edits in embryos were common, the ESbe2 hESC line with heterozygous editing at *HBG1* did not reveal any off-target activity of ABE **(Extended Data Fig. 7a-n)**.

As none of the Cas9-induced chromosome breaks identified in human embryos mapped to predicted off-target sites for the *PCSK9* sgRNA, we performed whole-genome sequencing (WGS) of genomic DNA from HUES64 hESCs treated in vitro with ABEmax RNP with *PCSK9* sgRNA, followed by Digenome-seq profiling^25–27^ to identify low-frequency off-target sites **(Extended Data Fig. 8a)**. We selected off-target sites based on i) the Levenshtein distance, a metric depending on the distance from the expected editing site^27,28^ from the *PCSK9* spacer sequence, or ii) the cleavage score, a measure of DSBs calculated by detecting concordant ends in forward and reverse reads as described previously for Digenome-seq analysis^25^ (**Supplementary Table 6**). Targeted deep sequencing of ABEmax-treated HUES64 cells did not reveal any off-target base editing at 25 sites selected by the Levenshtein distance (LD1-25) when compared to untreated controls **(Fig. 5e, Extended Data Fig. 8b, Supplementary Table 7)**. Similarly, we did not identify any evidence for off-target activity at the same sites upon analysis of a pooled WGA sample from 124 cells from 40 human embryos and in three edited hESC lines from *PCSK9* ABE-injected embryos (**Fig. 5f, Extended Data Fig. 8b, Supplementary Table 7**). We continued with assessing the base editing at a total of 121 genomic regions selected based on the cleavage score (CS1-121) in ABE-treated hESCs and untreated control **(Extended Data Fig. 8b-d)** as well as in human embryos and edited embryo-derived hESCs **(Fig. 5g, Extended Data Fig. 8b, e Supplementary Table 7)**. Through these analyses, we identified one off-target site (CS10) with a discernible base editing rate below 1% in human embryos. That same site was unedited in all three embryo-derived hESC lines **(Fig. 5g)**. In conclusion, ABE off-target editing is highly dependent on the sgRNA used; we identified a high frequency of off-target activity for *HBG1/2* sgRNA but minimal off-target activity for *PCSK9* sgRNA.

## DISCUSSION

In this study, we present that the ABE RNP injected into human zygotes, or at fertilization, is capable of efficient and precise modification of targeted genomic loci without detrimental chromosomal changes or large deletions. In addition, we found no adverse consequences of editing using ABE RNP for either preimplantation embryo development or genomic integrity of hESCs. Below we discuss two novel findings related to the basic biology of the human embryo as well as remaining limitations for the clinical use of base technology in early human embryos.

First, we report that the utilization of ABE is not associated with genotoxic effects, specifically large deletions and segmental chromosome aneuploidies. These genomic aberrations are often seen in blastomeres from human embryos edited with CRISPR/Cas9, suggesting that repair of DNA DSBs is compromised during early developmental stages^1^. In contrast to CRISPR/Cas9, ABE editors engage mismatch repair (MMR) and SSB repair^15^. DNA SSBs represent the most common DNA lesion arising from diverse sources. In human zygotes and early embryos, which undergo genome-wide DNA demethylation, SSBs and gaps may arise from the activity of the base excision repair pathway^29^, or during DNA replication^30^. The absence of large deletions and chromosomal changes at ABE-edited sites seen in our study suggests that nCas9-induced SSBs are not often processed into DNA DSBs and, therefore, genomic stability is maintained at target sites in human embryos. We did observe short insertions at a low frequency at *HBG1/2* loci in human blastomeres, and these insertions were adjacent to two target bases within the editing window. This indicates that a tandem of inosines in the template for MMR increases the error rate and mutagenic potential.

Second, our study revealed an unexpected mechanism for embryo arrest: exogenous mRNA. Injection of ABE RNP led to a significant improvement in the embryo’s developmental potential when compared to ABE mRNA injections. At the same time, our finding of mRNA toxicity points to a quality control mechanism sensing abnormal RNA in human embryos and resulting in early embryo arrest. Although used ABE mRNA was capped, modified with pseudouridine, and polyadenylated for improved stability and mimicking eukaryotic mRNA molecules, arrest invariably occurred. We show that the protein encoded is not the cause for the observed toxicity, as RNP allowed efficient development, enabling the derivation of homozygous and heterozygous edited hESC lines. Interestingly, upregulation of genes involved in mRNA recognition and belonging to interferon alpha and gamma responses was recently detected in human hematopoietic stem/progenitor cells treated with ABE or CBE mRNA, and improvement in mRNA design diminished this cellular sensing^20^. However, as there is essentially no transcription during the beginning of human preimplantation development, different mechanisms may apply. Interestingly, two recent studies reported development to the blastocyst stage only when ABE mRNA was applied at later stages of the development (e.g., eight-cell stage)^31,32^, suggesting the response to abnormal RNA may be developmentally regulated. Future studies should focus on the mechanisms of RNA sensing in human embryos and determining the physiological relevance to spontaneous embryo arrest.

Our work also highlights continued limitations of base editing in embryos. Mapping of Cas9-induced chromosome breaks and subsequent comparison with predicted off-target editing led to the identification of edited loci in human embryos beyond the intended target site. As ABE off-target editing activity was sgRNA dependent, guide testing and selection may avoid unintended editing. Furthermore, genetic mosaicism represents a yet unsolved obstacle for human embryo editing. Our sequencing analysis shows that in most cases, injection at the 1-cell stage results in mosaic embryos. Such an outcome prevents accurate genetic characterization at the blastocyst stage when a trophectoderm biopsy is commonly used to determine the genotype, albeit not the inner cell mass cells that form the fetus. Our study does not currently have sufficient data to comprehensively assess the frequency of mosaicism after ABE injections into MII oocytes at ICSI. To avoid embryo mosaicism, base editing before S-phase at ∼5-12 hours post fertilization will be needed.

In our study, the choice of editing targets, *HBG1/2* and *PCSK9*, was driven by the availability of normal human zygotes and previously tested sgRNAs, not therapeutic relevance to germline gene editing. Neither edit introduced here would be eligible for germline gene editing according to the latest report by the National Academies of Sciences, which recommends that the introduction of only naturally occurring and common genetic variants should be considered for heritable gene editing^33^. For most couples, preimplantation genetic testing for monogenic disorders (PGT-M) can be used to prevent diseases like sickle cell disease^34^. However, due to low live birth rates for PGT-M, genome editing may have merit in increasing the number of embryos available for implantation, thus raising the prospect of a genetically related healthy child^35,36^.

Furthermore, a recent study suggests that the incidence of numerous common disorders may be prevented through heritable polygenic editing (e.g., to prevent coronary heart disease through editing at *PCSK9*)^37^. Others have highlighted various ethical questions associated with heritable gene editing^38–41^. We expect that the data provided here will contribute to the conversations surrounding the risks and benefits of embryo editing. Notably, we show that the use of ABEs avoids the highly genotoxic effects seen after Cas9 targeting in preimplantation human embryos and enables normal development. Though this may be a step towards heritable editing, translation to a clinical context remains premature, with additional studies needed to address genetic mosaicism and the timing and uniformity of editing at on- and off-target sites.

## EXTENDED DATA FIGURES

**Extended Data Figure 1.**
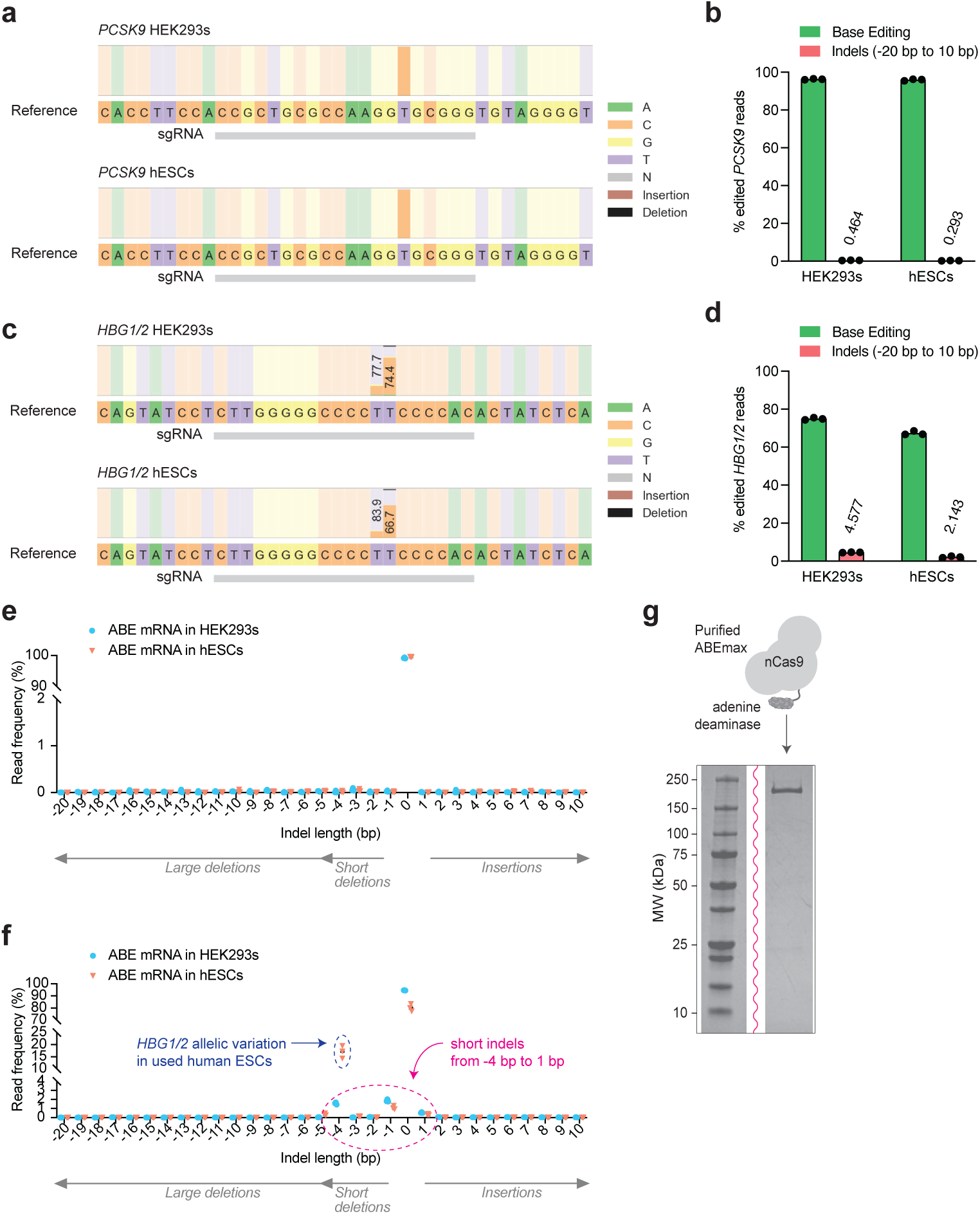
Analysis of adenine base editing efficiency and precision in human cells. **a, b,** Nucleotide percentage distribution around *PCSK9* sgRNA (**a**) and editing efficiency together with detected indels (analyzed length range -20 bp to 10 bp) (**b**) after adenine base editor (ABE) mRNA transfection of human embryonic kidney (HEK293T) cells or human embryonic stem cells (hESCs). **c, d,** Nucleotide percentage distribution around *HBG1/2* sgRNA (**c**) and editing efficiency together with detected indels (analyzed length range: -20 bp to 10 bp) (**d**) after ABE mRNA transfection of HEK293T cells or hESCs. **e, f,** NGS read frequency of individual indels within the indicated length range around *PCSK9* (**e**) and *HBG1/2* (**f**) target sites following ABE mRNA transfection of HEK293T cells and hESCs. In all panels, *n* = 3 replicates for both cell types. Data were analyzed, and panels (**a**) and (**c**) were prepared using Crispresso2^47^. **g,** Image showing sodium dodecyl sulfate–polyacrylamide gel electrophoresis (SDS-PAGE) analysis of the purified adenine base editor (ABE) protein, version ABEmax. nCas9, Cas9 nickase.

**Extended Data Figure 2.**
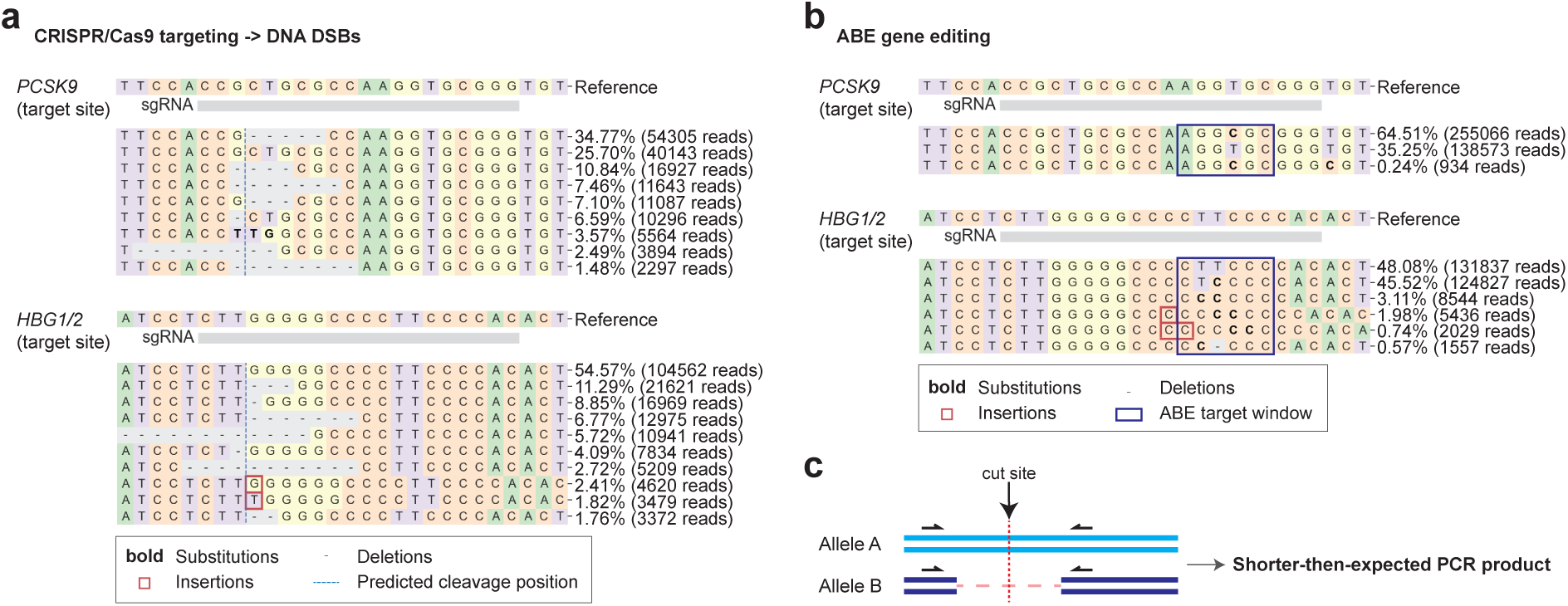
Comparison of on-target activity of CRISPR/Cas9 and adenine base editor in human embryos. **a, b,** Alleles frequency tables around *PCSK9* (upper panel) and *HBG1/2* (lower panel) sgRNAs after NGS using pooled WGA samples from CRISPR/Cas9-targeted (**a**) and ABE-edited (**b**) human embryos. *n* = 24 WGA blastomere samples were pooled for CRISPR/Cas9 NGS analysis, *n* = 124 WGA blastomere samples were pooled for NGS analysis of ABE with *PCSK9* sgRNA, and *n* = 27 WGA blastomere samples were pooled for NGS analysis of ABE with *HBG1/2* sgRNA. Data were analyzed, and panels (**a**) and (**b**) were prepared using Crispresso2^47^. **c,** Schematic showing large deletion on one allele around the target site of gene editor and the possibility of its detection via long-range PCR.

**Extended Data Figure 3.**
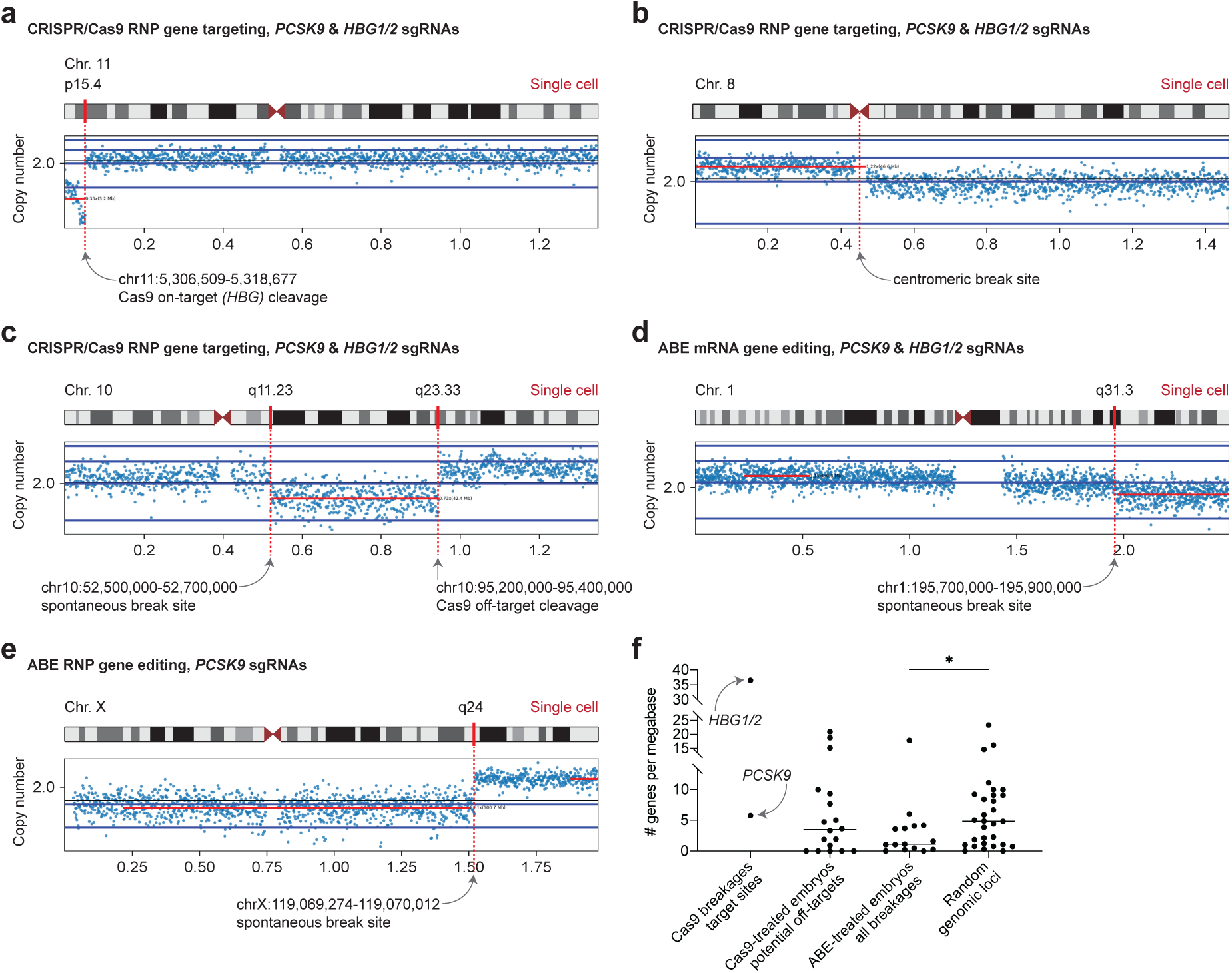
Chromosome breakage analysis in single cells from CRISPR/Cas9- or adenine base editor-injected human embryos. **a-e,** Copy number plots after SNP arrays of single blastomeres collected from human embryos injected at the zygote stage with CRISPR/Cas9 or ABE and *PCSK9 and HBG1/2* sgRNAs. Each break site was mapped visually and using SNP array data sets with heterozygosity calls. Dashed red lines indicate the position of the target site in copy number plots. Presented panels show chromosome break at the Cas9 target site *(HBG)* (**a**), break at the centromeric region (**b**), combination of spontaneous and Cas9-induced breaks (as determined by off-target analysis in this study) (**c**), and spontaneous breaks detected in ABE mRNA- or RNP-injected embryos (**d, e**). **f,** Gene density around: i) *PCSK9* and *HBG1/2* target sites; ii) candidate off-targets identified through chromosome breakage analysis (*n* = 18); iii) chromosome break sites in cells from ABE-injected embryos (*n* = 15); and iv) random genomic loci (*n* = 30) from a recent study^24^. Statistical significance established with a nonparametric *t*-test.

**Extended Data Figure 4.**
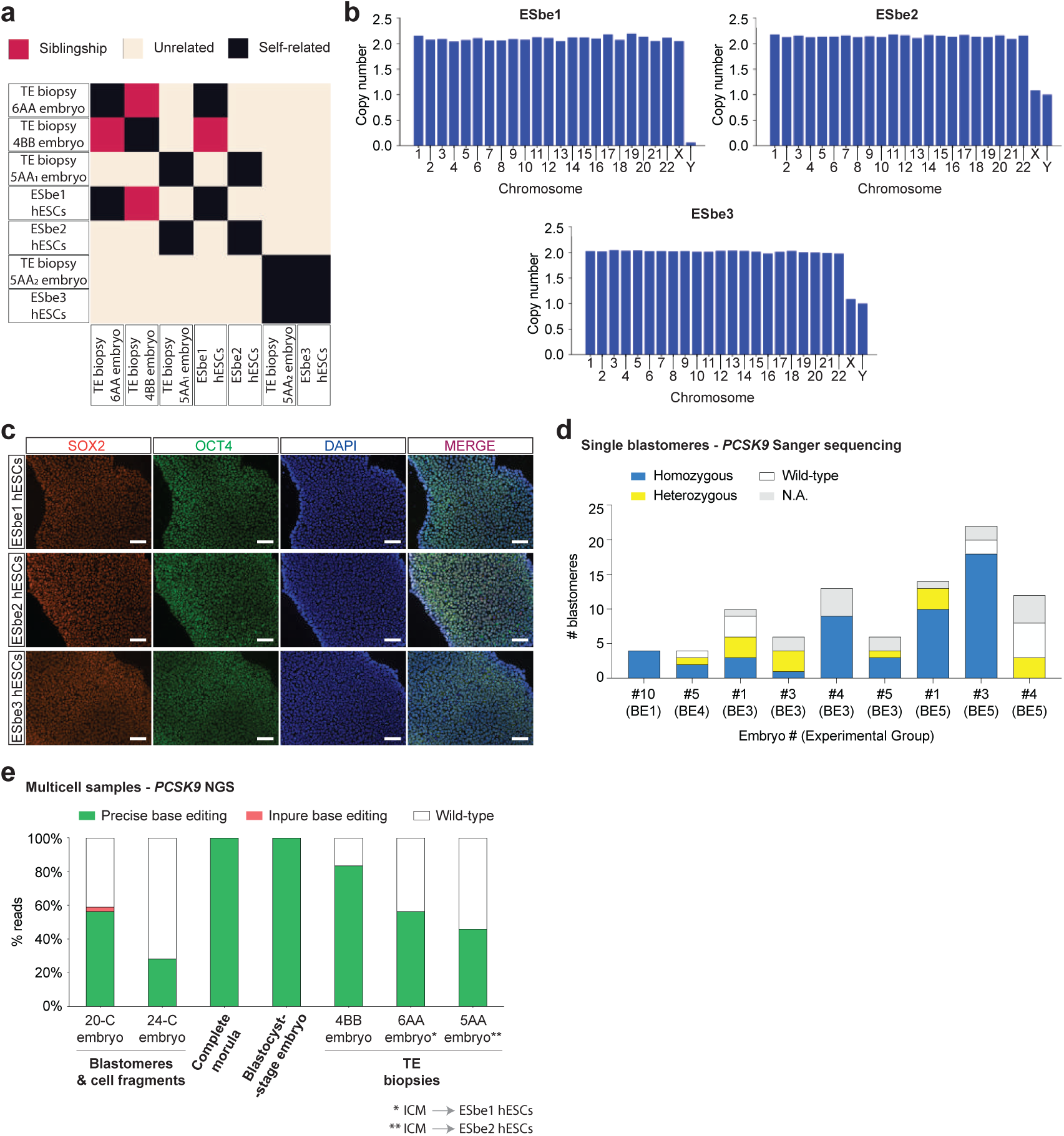
Development of adenine base-edited human embryos and characterization of derived base-edited embryonic stem cell lines. **a,** Heatmap generated based on SNP array data sets for selected trophectoderm (TE) and human embryonic stem cell (hESC) samples showing their genetic relatedness. **b,** Bar charts demonstrating that three hESC lines derived from base-edited human embryos are euploid. **c,** Immunostaining of three edited hESC lines derived from ABE RNP-injected human embryos showing expression of pluripotency markers SOX2 and OCT4. DNA stained by DAPI. Scale bars: 100 μm. **d,** Summary of Sanger sequencing analysis of genetic mosaicism at the *PCSK9* target site in human embryos injected with ABE at the zygote stage. Only the embryo #10 (experimental group BE1) is uniform, containing only cells edited on both *PCSK9* alleles. Embryo #4 (experimental group BE3) might also be uniform; however, 4/13 cells were not available for analysis. N.A., not available due to failed whole-genome amplification. **e,** Quantification after NGS analysis of adenine base editing at the *PCSK9* target site in multicell samples collected from human embryos at different stages of preimplantation development. Three samples show uniform editing: a complete morula- and blastocyst-stage embryo with 100% edited NGS reads (uniform homozygous editing) and the TE biopsy from a 5AA embryo with 50% edited and 50% unedited NGS reads (uniform heterozygous editing). The hESC line ESbe2 derived from the 5AA embryo shows heterozygous *PCSK9* editing (Fig. 4, panel h). The Gardner grading system was used to mark the quality of blastocysts. ICM, inner cell mass.

**Extended Data Figure 5.**
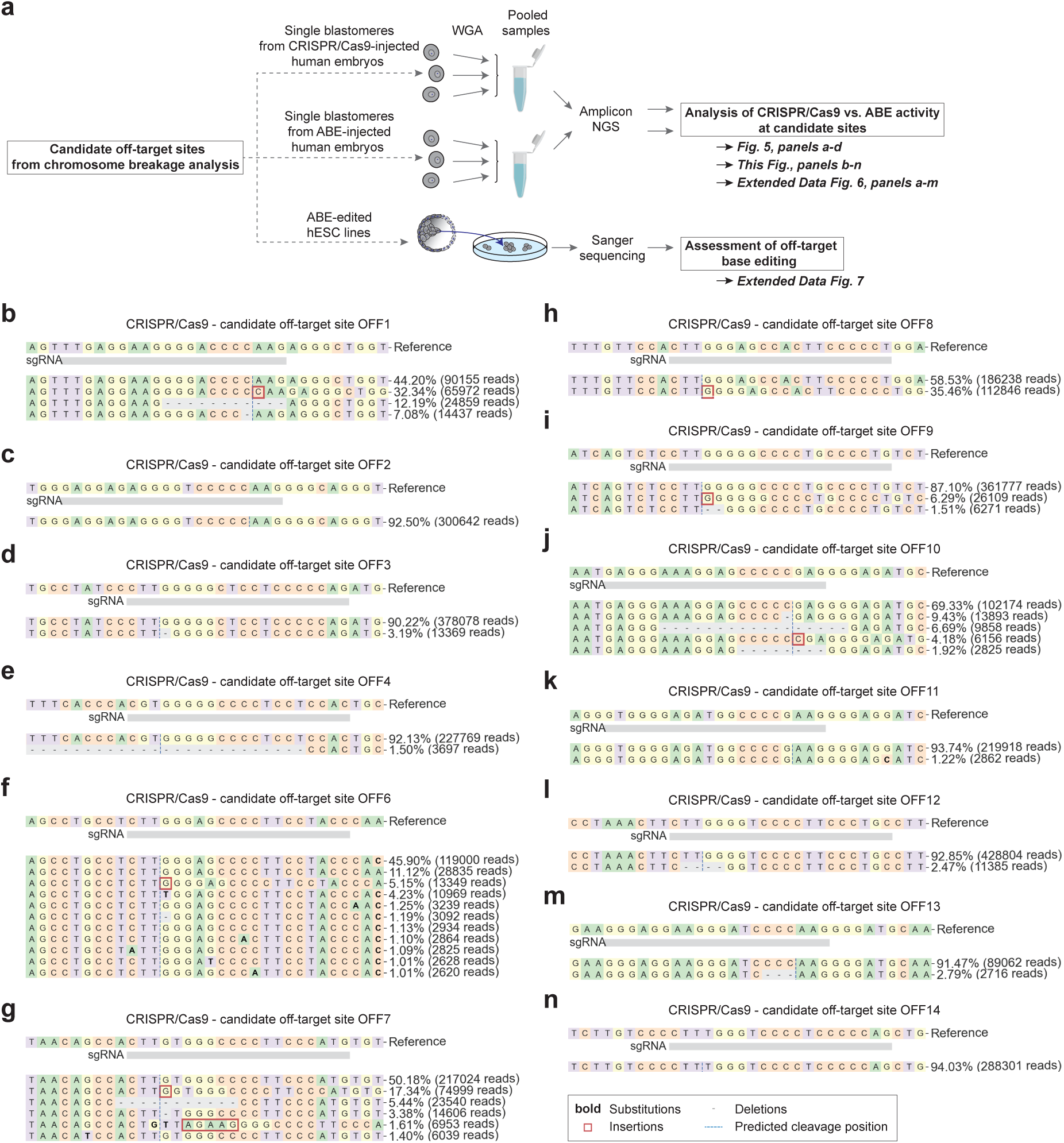
Cas9-induced indels at potential off-target sites identified through chromosome breakage analysis. **a,** Workflow of the assessment of ABE activity at candidate off-target sites OFF1-14 identified through chromosome breakage analysis in human embryos. Analysis of amplicon next-generation sequencing (NGS) was performed using pooled samples of whole-genome amplified (WGA) blastomeres collected from CRISPR- or ABE-injected embryos. Additionally, Sanger sequencing analysis was done using base-edited hESC line ESbe2 derived from a human embryo injected with ABE mRNA and *HBG1/2* sgRNA. **b-n,** Alleles frequency tables around 13 candidate off-targets after NGS using a pooled WGA sample from CRISPR/Cas9-injected human embryos. The result for candidate off-target site OFF5 is shown in Figure 5, panel a. *n* = 24 WGA blastomere samples were pooled for the analysis. Data were analyzed, and all panels were prepared using Crispresso2^47^. Protospacers of candidate off-targets are indicated by positions of the *HBG1/2* sgRNA (grey bar) below the reference DNA sequence in each panel.

**Extended Data Figure 6.**
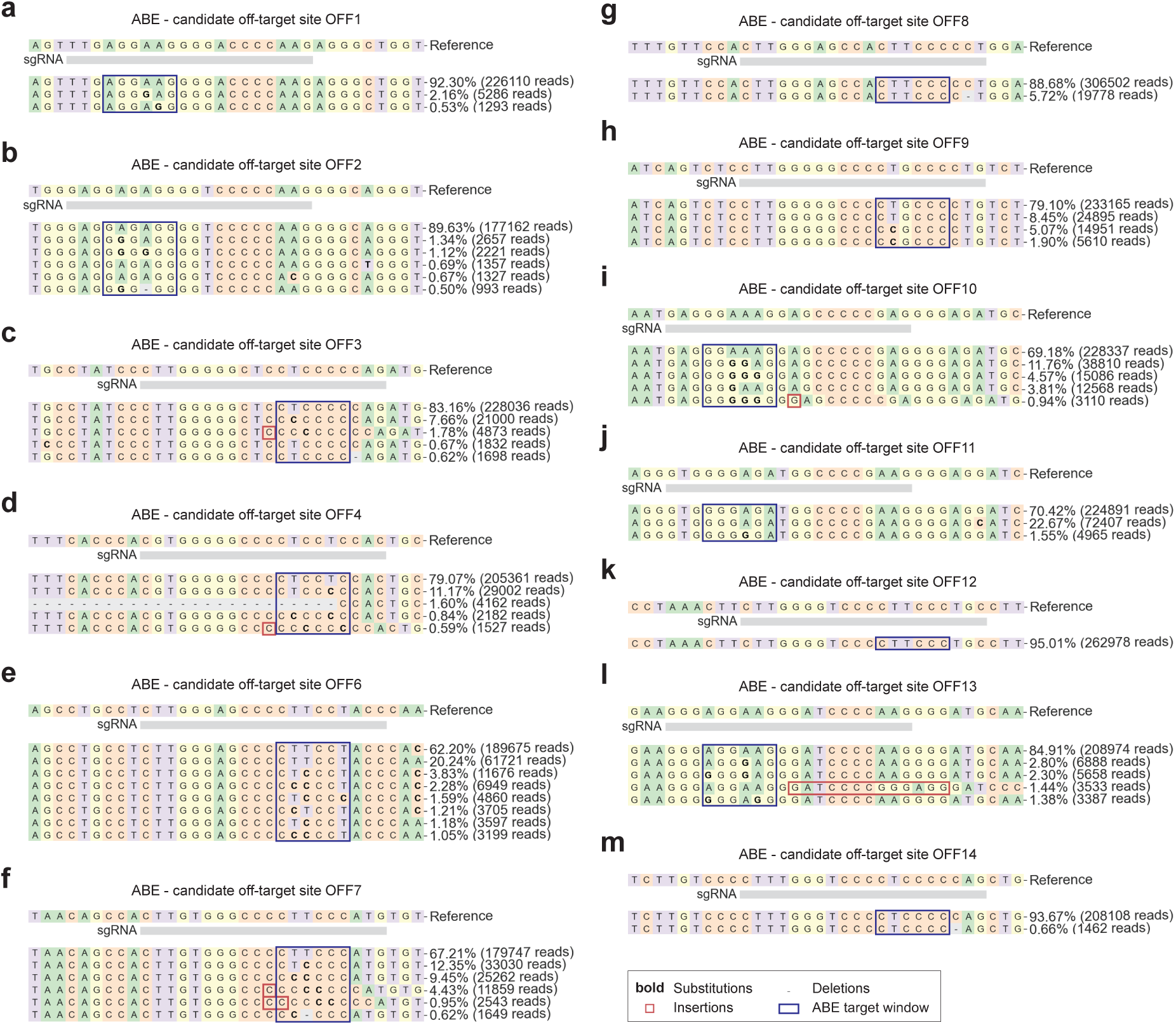
Base editing at potential off-target sites identified through chromosome breakage analysis. **a-m**, Alleles frequency tables around 13 candidate off-targets after NGS using a pooled WGA sample from ABE-injected human embryos. The result for candidate off-target site OFF5 is shown in Figure 5, panel b. *n* = 27 WGA blastomere samples were pooled for the analysis. Data were analyzed, and all panels were prepared using Crispresso2^47^. Protospacers of candidate off-targets are indicated by positions of the *HBG1/2* sgRNA (grey bar) below the reference DNA sequence in each panel.

**Extended Data Figure 7.**
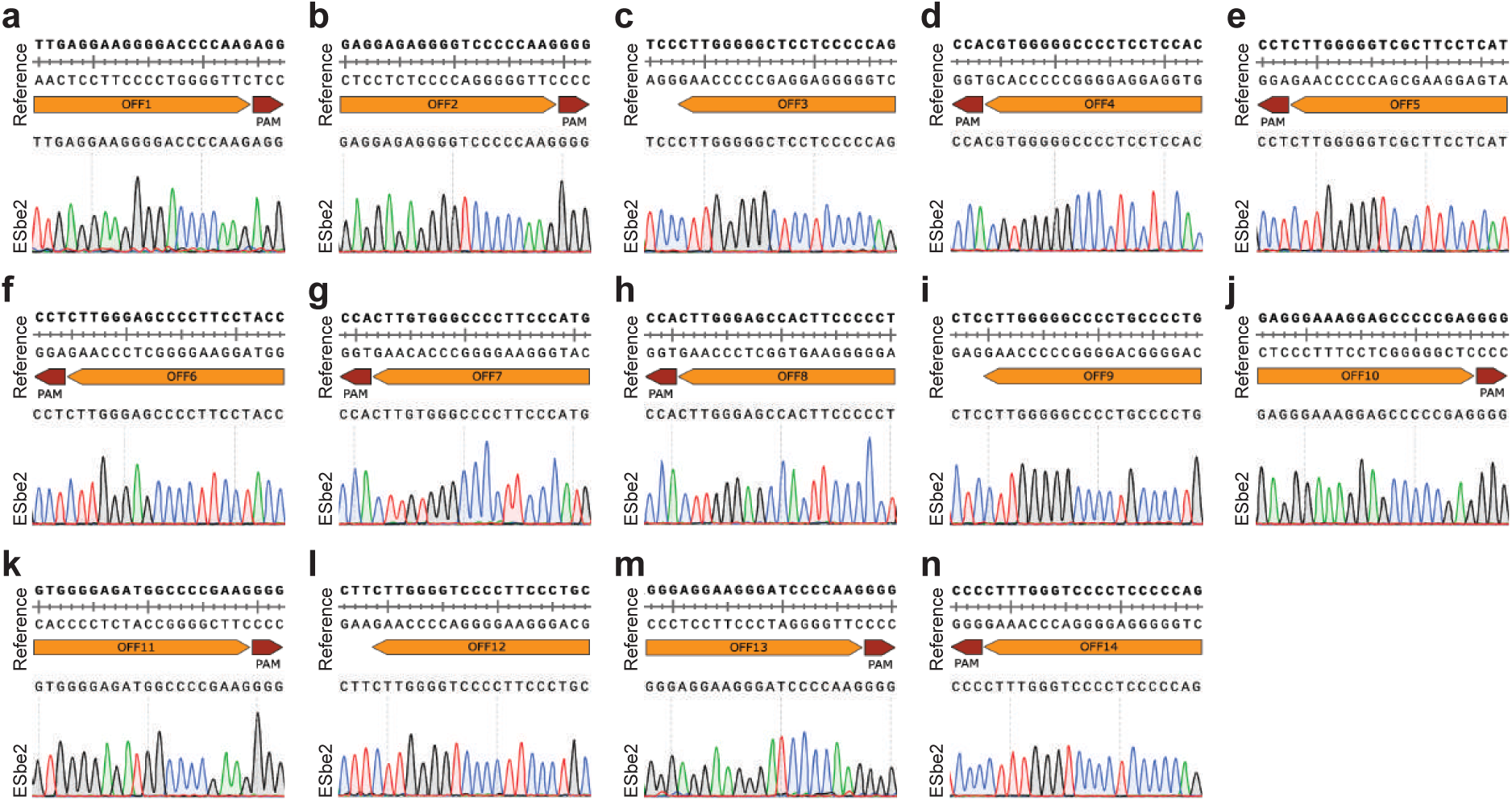
Undetectable off-target editing in human embryonic stem cells derived from a base-edited embryo at sites identified via chromosome breakage analysis. **a-n**, Sanger sequencing chromatograms used to analyze base editing at candidate off-target sites in ESbe2 hESCs derived from a human embryo injected with ABE mRNA and *HBG1/2* sgRNA at the zygote stage. All 14 off-target sites (OFF1-14) analyzed in this figure were identified through chromosome breakage analysis in human embryos.

**Extended Data Figure 8.**
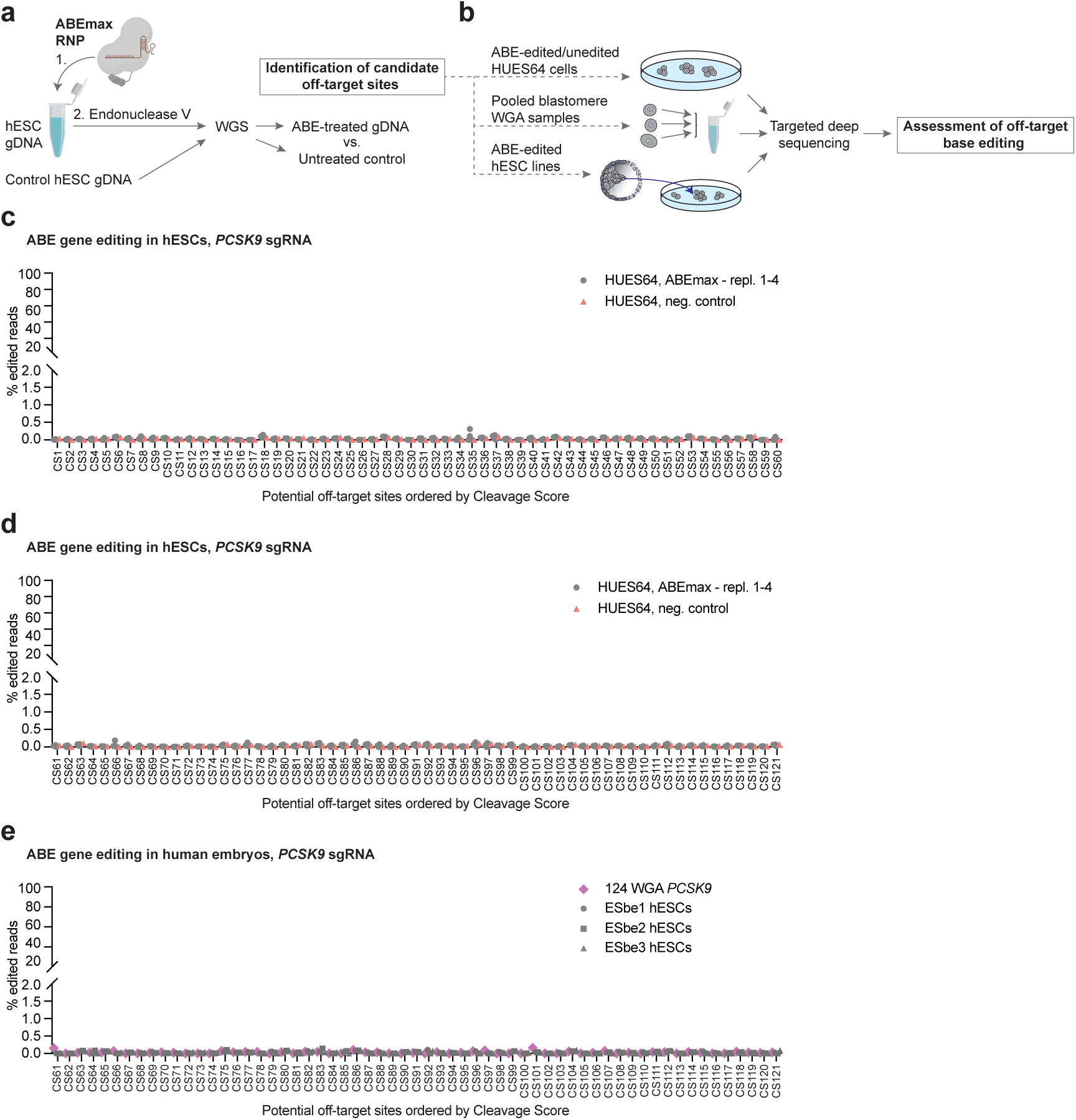
Undetectable off-target editing in human embryos and in human embryonic stem cells derived from base-edited embryos at sites identified via Digenome-seq. **a,** Workflow for identification of candidate ABE off-target sites. Genomic DNA (gDNA) isolated from HUES64 hESCs was incubated with ABEmax ribonucleoprotein (RNP), followed by treatment with endonuclease V and whole-genome sequencing (WGS). Candidate off-target sites were selected after Digenome-seq analysis and comparison with an untreated HUES64 hESC control sample. **b,** Assessment of ABE activity at candidate off-target sites via targeted deep sequencing in: i) ABE-treated HUES64 hESCs; ii) pooled WGA samples from base-edited embryos; and iii) hESC lines derived from ABE-injected embryos. **c, d,** ABE gene editing efficiency at potential off-targets (ordered by cleavage score; CL1-121) in transfected HUES64 hESCs (*n* = 4) and untreated control. Panel (**c**) shows analysis of candidate off-targets CL1-60, and panel (**d**) shows analysis of candidate off-targets CL61-121. **e,** ABE gene editing efficiency at potential off-targets (ordered by cleavage score; CL61-121) in human embryos and three hESC lines derived from base-edited embryos. *n* = 124 WGA blastomere samples were pooled for the NGS analysis.

## EXTENDED DATA TABLES

**Extended Data Table 1.**
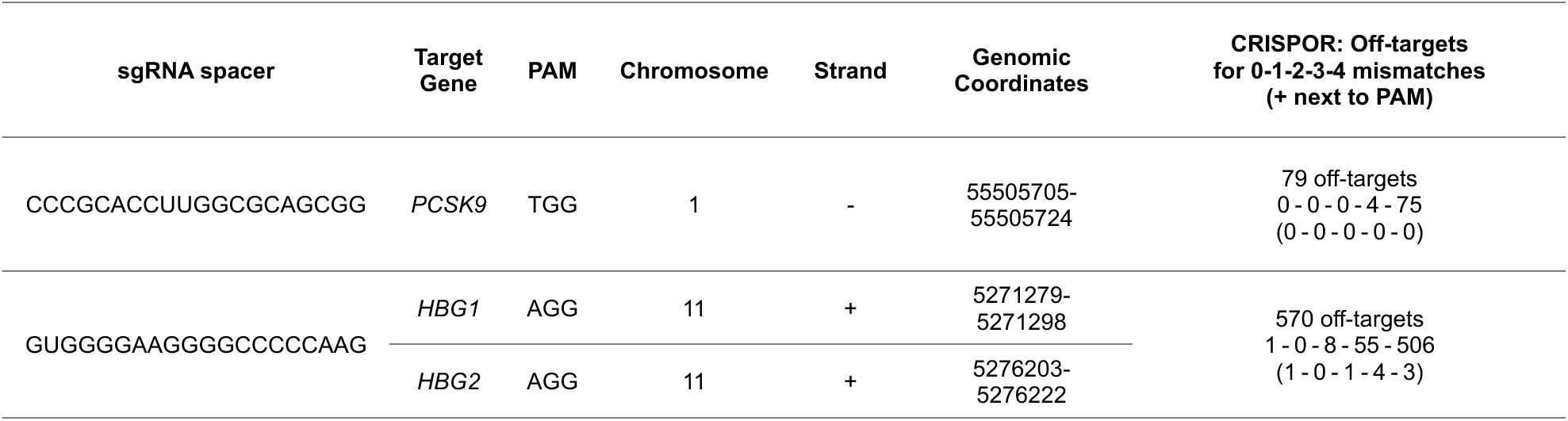
Guide RNAs and genomic targets.

**Extended Data Table 2.**
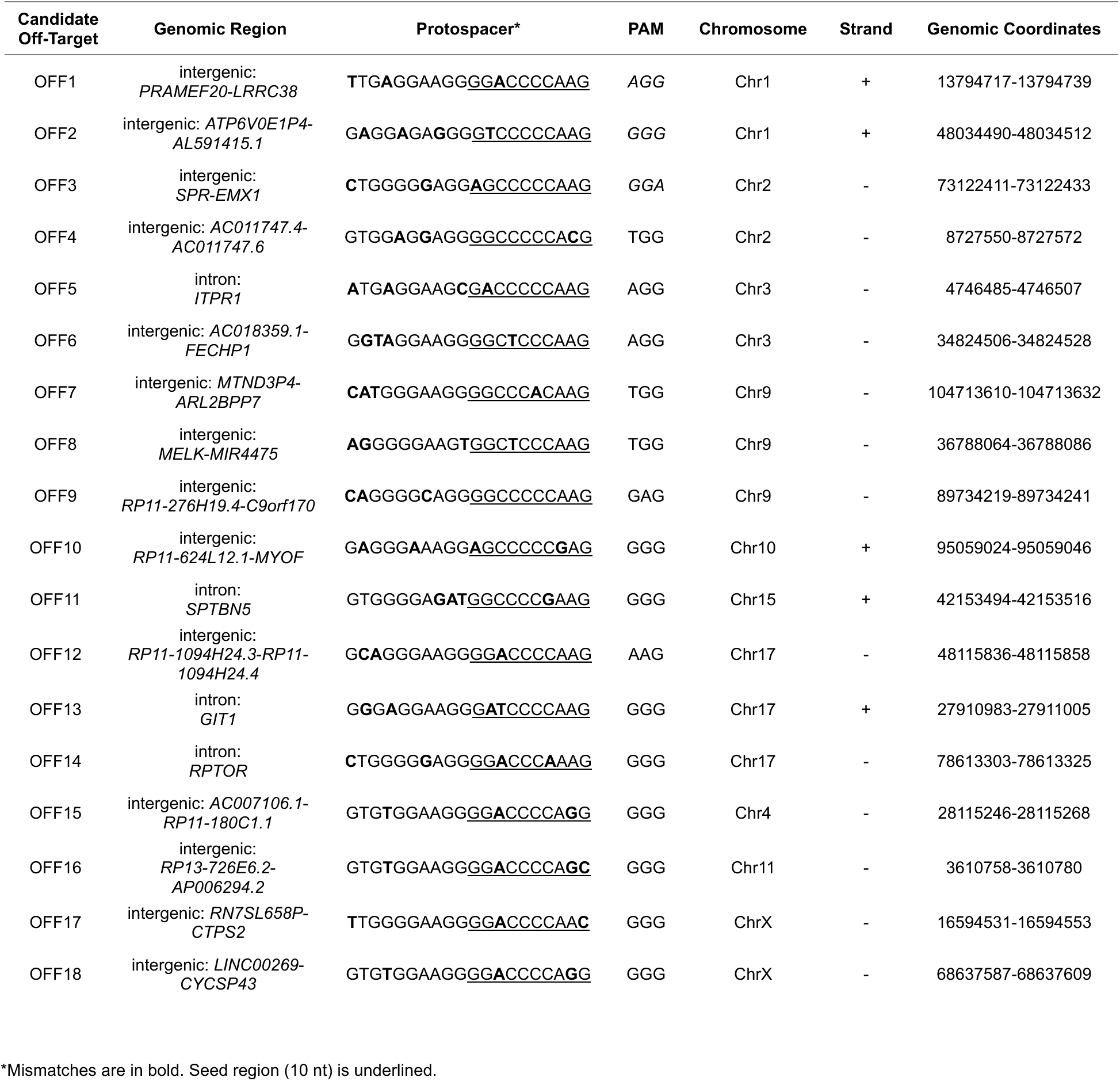
Candidate off-target sites identified by chromosome breakage analysis.

**Extended Data Table 3.** Developmental potential of human embryos after ABE injections.

|  | Embryo #<br>within each cohort | ABE mRNA | ABE RNP |  |
| --- | --- | --- | --- | --- |
|  |  | 2PN zygotes | 2PN zygotes | MII oocytes |
| Cell number | 1 | 4 | 100 | 100 |
|  | 2 | 4 | 100 | 100 |
|  | 3 | 3 | 24 | 100 |
|  | 4 | 2 | 20 | 100 |
|  | 5 | 2 | 14 | 9 |
|  | 6 | 2 | 13 | 8 |
|  | 7 | 2 | 12 | 8 |
|  | 8 | 2 | 10 | 7 |
|  | 9 | 2 | 6 | 5 |
|  | 10 | 2 | 6 | 4 |
|  | 11 | 2 | 6 | 3 |
|  | 12 | 2 | 4 | 1 |
|  | 13 | 2 | 3 |  |
|  | 14 | 1 |  |  |
|  | 15 | 1 |  |  |
|  | 16 | 1 |  |  |
|  | 17 | 1 |  |  |
|  | 18 | 1 |  |  |
|  | 19 | 1 |  |  |

Number of cells per each human embryo is listed. Embryos are sorted according to their developmental potential (from largest to smallest cell number).
Blastocyst are estimated to consist of 100 cells (a lower estimate).

## METHODS

### Human Subjects Research

The work with human gametes, embryos, and embryonic stem cells was reviewed and approved by the Columbia University Human Embryonic and Human Pluripotent Stem Cell Research Committee and the Institutional Review Board.

### Gamete and Embryo Donation

MII oocytes were obtained from anonymous donors for the purpose of fertilization with consent for use in research. 2PN zygotes were donated by patients who decided that their treatment in the fertility center was complete and that the embryos were no longer needed for the purpose of reproduction. Donation to research was chosen in lieu of destruction. All embryos were de-identified prior to use in research.

### Purification of Adenine Base Editor

Protein expression and purification were performed as described by Jang and colleagues^42^. Briefly, HEK293T cells were transiently transfected with pEX-FlagR-ABEmax (NGG PAM) plasmid designed to express the base editor as a fusion protein with a 10xHis-FLAG-tag at the N-terminus and an mCherry-10xHis-tag at the C-terminus. Cells were transfected at a density of 7 × 10^5^ cells/ml using 25-kDa linear polyethyleneimine (Polysciences), and dimethyl sulfoxide (Amresco) was added immediately after the transfection to a final concentration of 1%. The temperature in the incubator was then lowered to 33°C. Two days after transfection, tryptone (Amresco) was added to a final concentration of 0.5%. The cells were harvested four days after transfection, resuspended in a lysis buffer [20-mM Tris-HCl (pH 7.5), 1-M NaCl, 2-mM β-mercaptoethanol, and 20% glycerol], and lysed by sonication. The sonicated sample was centrifuged, and the supernatant was loaded on an Ni-NTA column (Qiagen). The column was washed with buffer A [20-mM Tris-HCl (pH 7.5), 500-mM NaCl, 2-mM β-mercaptoethanol, and 20% glycerol] supplemented with 40-mM imidazole (MilliporeSigma), and bound proteins were eluted with buffer A supplemented with 200-mM imidazole. Eluted proteins were treated with tobacco etch virus (TEV) protease and human rhinovirus (HRV) 3C protease overnight to expose the FLAG-tag at the N-terminus and remove the C-terminal mCherry-10xHis-tag, respectively. The sample was then mixed with α-FLAG M1 agarose resin (MilliporeSigma) in the presence of 5-mM CaCl_2_ and rotated slowly at 4°C for 1 hour. After a wash with buffer A supplemented with 1-mM CaCl_2_, the bound proteins were eluted with buffer A supplemented with 5-mM ethylene glycol-bis(2-aminoethyl ether)-N,N,N′,N′-tetraacetic acid (EGTA). The eluted FLAG-ABEmax was concentrated and further purified using a HiLoad 16/600 Superdex 200-pg column equilibrated with a storage buffer [20-mM Tris-HCl (pH 7.5), 500-mM NaCl, 1-mM β-mercaptoethanol, and 20% glycerol]. The peak fraction was concentrated to a final concentration of ∼10 µg/µl, flash-frozen in liquid nitrogen, and stored at −80°C.

### In Vitro Transcription

A plasmid encoding ABE8.8-m was obtained from Addgene (plasmid no. 136294). Base editor ABE8.8.-m was PCR amplified with primers to include the T7 promoter and install a 120-nt poly(A) tail to the 3’ untranslated region (**Supplementary Table 8)**. The PCR product was purified with the QIAquick PCR Purification Kit (QIAGEN) and then used as a template for in vitro transcription (IVT) using the HiScribe T7 High-Yield RNA Synthesis Kit (New England Biolabs). Co-transcriptional capping with CleanCap AG (TriLink BioTechnologies; Cat. no. N-7113) and full replacement of uridine-5′-triphosphates for N1-Methylpseudouridine-5’-Triphosphates (TriLink BioTechnologies; Cat. no. N-1081) was performed during the IVT. Synthesized RNA was purified by precipitation with 2.5-M LiCl (Thermo Fisher Scientific), followed by two wash steps in 70% ethanol and dissolving in nuclease-free water (Integrated DNA Technologies). The final concentration was measured on a NanoDrop 1000 Spectrophotometer (Thermo Fisher Scientific), aliquoted, and stored at −80°C.

### Ribonucleoprotein Assembly

Both *PCSK9* and *HBG1/2* sgRNAs (**Extended Data Table 1**) were purchased from Integrated DNA Technologies (IDT) and dissolved in IDTE buffer (10-mM Tris, 0.1-mM EDTA, pH 7.5) to prepare 100 µM stocks. Alt-R S.p. Cas9 Nuclease V3 (10 µg/µl) was obtained from IDT. For CRISPR/Cas9 RNP assembly, 3 μl of injection buffer (5-mM Tris-HCl, 0.1-mM EDTA, pH 7.8), 2 μl of Alt-R S.p. Cas9 Nuclease V3, and 1.5 μl of 100-µM sgRNA were combined and incubated at room temperature for 15 minutes, followed by the addition of 93.5 μl of injection buffer. Diluted Cas9 RNP was centrifuged at 16,000 x g for 2 minutes prior to the injections to human oocytes or embryos.

For base editing, ABE RNP was prepared similarly as described above, only substituting Cas9 nuclease with an equimolar amount of purified ABEmax protein (12.25 µg/µl). If multiplexed targeting of *PCSK9* and *HBG1/2* was performed, identical amounts of solutions of RNP with the two sgRNAs were combined, mixed, and centrifuged prior to the injections.

### Intracytoplasmic Sperm Injection

Prior to intracytoplasmic sperm injection (ICSI), cryopreserved sperm was thawed by 10-minute incubation at room temperature and transferred to a 15-ml conical centrifuge tube. Sperm Washing Medium (FUJIFILM Irvine Scientific; Cat. no. 9983) was added dropwise to a final volume of 6 ml. The sample was centrifuged at 300x g for 15 minutes. The supernatant was discarded, and an additional wash was performed. The sperm pellet was resuspended in the wash medium and analyzed for viability and motility.

Manipulation dishes contained a droplet with 10% polyvinylpyrrolidone solution with human serum albumin (FUJIFILM Irvine Scientific; Catalog ID 90123), a droplet with ABEmax RNP in injection buffer, and a droplet of global total with HEPES (LifeGlobal Group; Cat. no. LGTH-050). Sperm was mixed with the PVP solution, and a single motile sperm was immobilized by pressing its tail with the ICSI micropipette. The selected sperm was then picked, ejected in the RNP droplet, and picked up again for injections. After all manipulation, injected cells were cultured in dishes containing global total (LifeGlobal Group; Cat. no. LGGT-100) and placed in an incubator at 37°C and 5.5% CO_2_ and cultured until collection. The pronucleus formation was confirmed on day 1 after ICSI.

### Embryo Manipulation and Microinjection

All human oocytes and embryos were manipulated as previously described. For thawing of cryopreserved human 2PN zygotes, cryo straws were held at room temperature air until liquid nitrogen dissipated, followed by thawing in a water bath at 37°C for 3 minutes. Cryo straws were wiped and cut at the ends, and the content of each cryo straw was pushed into a culture dish. Embryos were located and subsequently moved into a new dish containing global total with HEPES (LifeGlobal Group; Cat. no. LGTH-050) for 5 minutes. Embryos were then placed in an incubator in global total (LifeGlobal Group; Cat. no. LGGT-100) for an additional 10-15 minutes before injections were performed.

Injection needles were made from borosilicate glass capillaries (Sutter Instrument; Cat. no. B100-75-10) using a P-1000 Micropipette Puller (Sutter Instrument) and a program with the following parameters: Heat 578, Pull 0, Velocity 200, Delay 1, and Pressure 500. Embryos were placed on the heated stage of an inverted Olympus IX73 microscope in droplets of global total with HEPES covered by mineral oil and injected with 50-200 pl of a mixture of in vitro synthesized mRNA with synthetic sgRNA, or the RNP. After injections, embryos in global total were placed in an incubator at 37°C and 5.5% CO_2_ and cultured until collection.

### Derivation of Embryonic Stem Cells

Embryonic stem cells were derived as previously described^43^. Briefly, a trophectoderm biopsy of 5-10 cells was taken using laser-assisted micromanipulation. Two-thirds of the trophectoderm (excluding the trophectoderm near the ICM) was ablated using laser (Hamilton Thorne) pulses (300 µs at 100% intensity)^44^, and the remaining ICM was plated on an MEF feeder layer with ESC derivation medium [KnockOut DMEM (Thermo Fisher Scientific; Cat. no. 10829018), Gibco GlutaMAX Supplement (Thermo Fisher Scientific; Cat. no. 35050061; dilution 1:100), Gibco MEM Non-Essential Amino Acids (Thermo Fisher Scientific; Cat. no. 11140050; dilution 1:100), 10,000 U/ml Penicillin-Streptomycin solution (Thermo Fisher Scientific; Cat. no. 11140050; final concentration 100 U/ml) and Gibco 2-Mercaptoethanol (Thermo Fisher Scientific; Cat. no. 21985023; dilution 1:1,000)] supplemented with 5% HyClone FBS (Cytiva) and the following small molecules (Tocris Bioscience): Chroman 1 (Cat. No. 7163; dilution 1:10,000), Emricasan (Cat. No. 7310; dilution 1:10,000), Polyamine Supplement x1000 (Cat. No. 7739; dilution 1:1,000), and trans-ISRIB (Cat. No. 5284; dilution 1:10,000). Attachment was monitored on day 2 after plating, with laser removal of excess trophectoderm growth. Half medium was changed every second day. After the emergence of the outgrowth, the ROCK inhibitor was removed, and after ∼2 weeks, when ∼0.1-0.5 cm in diameter, a glass microcapillary was used to cut the colony into smaller pieces and plated on a StemFlex-based culture system for passaging and cryopreservation, as well as isolation of gDNA.

### Genome-wide Single Nucleotide Polymorphism Array

Embryos were treated with EmbryoMax Acidic Tyrode’s solution (EMD Millipore; Cat. no. MR-004-D) to remove the zona pellucida and placed in droplets of PGD Biopsy Medium (LifeGlobal; Cat. no. LPGG-020) covered by mineral oil. Embryos were dissected to individual cells using a Narishige micromanipulator equipped with two fine capillaries to pry individual cells apart. Each cell was then placed in 2 ul of PBS for WGA, followed by SNP array analysis and genotyping. Genome amplifications were performed at Genomic Prediction’s Clinical Laboratory using ePGT amplification protocol^45^. Amplified DNA was processed for Axiom GeneTitan and UK Biobank Axiom Arrays (Thermo Fisher Scientific) according to the manufacturer’s protocol. Copy number and genotyping analyses were performed using gSUITE software (Genomic Prediction).

### Chromosome Break Site Mapping

Identification of endogenous break sites was done using copy number plots and by visual evaluation of loss of heterozygosity. Genomic coordinates (Homo sapiens genome assembly GRCh37/hg19 was used in this study) for each chromosome breakage were determined through analysis of SNP array data sets using both copy number signals and heterozygosity calls. The accuracy of mapping is between 10 and 200 kb. For the comparison of gene densities around the chromosome breakages, a data set for random genomic loci was used from recent study^24^ (**Supplementary Table 3**). For the identification of potential off-target sites, genomic coordinates of chromosome breakages in cells isolated from Cas9-injected embryos were compared with in silico-predicted (CRISPOR^46^; http://crispor.org/) off-targets for used *PCSK9* and *HBG1/2* sgRNAs **(Supplementary Table 9).**

### Genotyping by Sanger Sequencing and Next-Generation Sequencing

Amplified DNA samples were diluted in nuclease-free water and used for PCR amplifications by AmpliTaq Gold 360 DNA Polymerase (Thermo Fisher Scientific). PCR primers are listed in **Supplementary Table 8**. PCR amplicons were visually inspected on agarose gel prior to Sanger sequencing or NGS at GENEWIZ (Azenta Life Sciences). Sanger sequencing traces were visualized and analyzed using Benchling. For NGS, PCR products were purified via sodium acetate precipitation or using the QIAquick PCR Purification Kit (QIAGEN), followed by submission for Amplicon-EZ sequencing at GENEWIZ. Sequencing results were received in the form of raw reads and fastq files, which were analyzed using CRISPResso2^47^. For evaluation of ABE activity at candidate off-targets identified through chromosome breakage analysis, all 14 sites (OFF1-14) that do not exhibit sequence variability were analyzed by NGS using pooled blastomere WGA samples from CRISPR- or ABE-injected embryos. Four potential off-target sites (OFF15-18) with alternative sequences that differ from the reference genome were not included in this analysis. In experiments involving WGA samples from a small number of cells, we set a read frequency threshold during the NGS analysis to avoid misinterpretation of mutations introduced during DNA amplifications and/or sequencing errors. This applied to NGS analyses of multicell embryo biopsies (threshold 1%; **Extended Data Fig. 4e**), pooled WGA sample of cells from Cas9-injected embryos (threshold 0.5%; **Fig. 5a, Extended Data Fig. 5b-n**), and pooled WGA sample of cells from ABE-injected embryos (threshold 0.5%; **Fig. 5b, Extended Data Fig. 6a-m**).

### Cell Culture Conditions for Human Cells

HEK293T cells were cultured in Dulbecco’s Eagle Modified Medium (DMEM) with GlutaMAX (Thermo Fisher Scientific; Cat. no. 10566016) supplemented with 10% fetal bovine serum (FBS) (Sigma-Aldrich). The cells were cultured at 37°C in a humidified incubator with 5% CO_2_ and passaged every 3-4 days at confluency below 80%. hESCs were cryopreserved in a solution of freezing media containing 40% fetal bovine albumin (FBS) and 10% dimethyl sulfoxide (Sigma-Aldrich).

hESCs were cultured in Gibco StemFlex medium (Thermo Fisher Scientific; Cat. no. A3349401) on Geltrex Matrix (Thermo Fisher Scientific; Cat. no. A1413301)-coated cell culture plates. The cells were cultured at 37°C in a humidified incubator with 5% CO_2_, and passaged every 5-7 days (at approximately 70% confluency) at a ratio 1:10. For passaging, TrypLE Express (Thermo Fisher Scientific; Cat no. 12605010) was used to dissociate the cells into small clusters, which were then plated in a medium containing 10-uM ROCK inhibitor Y-27632 (Sigma-Aldrich; Cat. no. SCM075). Fresh medium without ROCK inhibitor was added 24-48 hours after passaging. hESCs were cryopreserved in a solution of freezing media containing 40% fetal bovine albumin (FBS; GeminiBio; Cat. no. 900-108) and 10% DMSO (Sigma-Aldrich; Cat. no. D2650).

### Nucleofections of Human Cells

All cells were cultured as described in the previous section. Before nucleofection, hESCs cells were dissociated with TrypLE Express (Thermo Fisher Scientific; Cat no. 12605010), pipetted to fragment the colonies into small clusters consisting of 2-3 cells, and counted using the Countess 3 Automated Cell Counter (Thermo Fisher Scientific). 4.5×10^5^ cells were resuspended in 20 μl nucleofection solution with ABEmax RNP [assembled using 1.8 µl ABEmax (12.25 µg/µl) and 1.2 µl 100-uM sgRNA] and nucleofected using P3 Primary Cell 4D-Nucleofector X Kit (Lonza; Cat. No. V4XP-3032) and Lonza 4D-Nucleofector System (program CA-137) according to the manufacturer’s protocol. Cells were subsequently plated into new dishes with StemFlex medium supplemented with 10-uM ROCK inhibitor Y-27632 (Sigma-Aldrich; Cat. no. SCM075) and 100 μg/ml of Normocin (Invivogen; Cat. No. ant-nr-1). The medium was changed one day post-nucleofection to withdraw the ROCK inhibitor and antibiotics. Three days after transfection, cells were harvested, and genomic DNA was isolated using the DNeasy Blood and Tissue Kit (QIAGEN; Cat. No. 69504) for PCR amplification of targeted loci and sequencing analysis by NGS.

For nucleofections using mRNA, hESCs were cultured, harvested, and nucleofected following the same procedure as described above, except with replacing ABE RNP with 4 μg in vitro synthesized ABE8.8-m mRNA combined with 150 pmol synthetic sgRNA. Identical amounts of mRNAs were used also for nucleofections of HEK293T cells with differences in culture conditions related to these cells and the use of SF Cell Line 4D-Nucleofector X Kit S (Lonza; Cat. no. V4XC-2032). In **Extended Data Fig. 1**, panels (**a-f**) the used hESC line was derived in the Egli laboratory, and the cells carry a heterozygous mutation (rs758109813) at the *EYS* locus^1^.

### EndoV-coupled Digenome-seq for In Vitro Off-target Profiling of ABEmax

To evaluate off-target activity of ABEmax in vitro, ribonucleoprotein (RNP) complexes were assembled by incubating 465 pmol of in vitro transcribed sgRNA (synthesized via T7 in vitro transcription) with 155 pmol of ABEmax protein at room temperature for 30 minutes. The RNP complex was then incubated with 10 µg of genomic DNA (HUES64) in 2×BF buffer (Biosesang) at 37°C for 16 hours.

To remove residual sgRNA, 4 µL of RNase A (100 mg/mL) (QIAGEN) was added and incubated at 37°C for 15 minutes. The reaction mixture was then purified using the Gel and PCR Clean-up Kit (MACHEREY-NAGEL).

Subsequently, 3 µg of the purified genomic DNA was treated with 8 units of Endonuclease V (New England Biolabs) in NEBuffer 4 at 37°C for 2 hours. The final reaction was again purified using the Gel and PCR Clean-up Kit (MACHEREY-NAGEL), and the resulting DNA was subjected to WGS using the Illumina NovaSeqX platform. As a negative control, genomic DNA that had not been treated with ABEmax was processed in parallel and subjected to WGS.

### Off-target Site Identification via Digenome-seq

Potential off-target sites were identified by analyzing the WGS data from ABE-treated HUES64 hESCs and untreated control HUES64 hESCs samples using the Digenome-seq web tool (rgenome.net). Candidate sites were first filtered to include only those with a Levenshtein distance of 10 or less from the on-target sgRNA spacer sequence targeting *PCSK9*. Independently, the top 50 sites with the highest cleavage scores, as determined by the Digenome-seq tool^25^, were also selected. The union of these two sets was subjected to further analysis (**Supplementary Table 6**).

Sites that showed similar read patterns between the negative control and ABE-treated samples or that exhibited an abnormally high indel frequency were excluded from downstream analysis based on manual inspection using the Integrative Genomics Viewer (IGV).

### Targeted Deep Sequencing and A-to-T Conversion Analysis

For each selected site, targeted deep sequencing was performed to quantify A-to-T conversion rates in the ABE-treated and control samples. Primers for each region were designed using Primer-BLAST (https://www.ncbi.nlm.nih.gov/tools/primer-blast), and amplification of the targeted regions was carried out using KOD Multi & Epi DNA polymerase (TOYOBO) following the manufacturer’s protocol. Primers are listed in **Supplementary Table 8.**

The prepared libraries were sequenced using paired-end reads on the Illumina MiniSeq platform with the MiniSeq High Output Reagent Kit (300 cycles). The resulting FASTQ files were analyzed using custom scripts written in Python 3 to calculate base conversion frequencies (https://github.com/BaeLab/off-target-deepseq).

### Western Blot

The cultured hESC lines ESbe3 and HUES64 were homogenized in a protein lysis buffer containing protease inhibitors using the IntactProtein Cell-Tissue Lysis Kit (GenulN Biotech; Cat. no. 415). The protein concentration was determined using a BCA assay (Thermo Fisher Scientific; Cat. no. 23225). The equal amounts of protein extracts (50 μg) were loaded for polyacrylamide gel electrophoresis and subsequently transferred to polyvinylidene fluoride (PVDF) membranes. The membranes were blocked for 1 hour with 5% non-fat milk in Tris-buffered saline (20-mM Tris, 150-mM NaCl) and 0.1% (w/v) Tween 20 (TBST) and then incubated with primary antibodies for PCSK9 (Invitrogen; Cat. no. MA5-32843; dilution 1:500) and GAPDH (Proteintech; Cat. no. HRP-60004; dilution 1:2,000) at 4°C overnight. The protein bands were detected with HRP-conjugated secondary antibodies (Cytiva; Cat. no. NA934 and NXA931; dilution 1:5,000) and visualized by enhanced chemiluminescence (ECL) using Pierce ECL Western Blotting Substrate (Thermo Fisher Scientific; Cat. no. 32106).

### Immunostaining

Human embryonic stem cells (lines ESbe1, ESbe2, and ESbe3) were fixed by incubation in 4% (v/v) paraformaldehyde in phosphate-buffered saline (PBS) for 20 minutes at room temperature (RT), followed by rinsing three times with PBS. Cells were permeabilized by incubation in 0.1% Triton X-100 in PBS for 15 minutes at RT and washed three times with PBS. Cells were blocked in 5% bovine serum albumin (BSA) in PBS for 1 hour at RT. The following primary antibodies were diluted in 1% BSA in PBS and applied at 4°C for 16 hours: mouse monoclonal anti-OCT4 (Santa Cruz Biotechnology; Cat. no. sc-5279; dilution 1:200) and rabbit monoclonal anti-SOX2 (Cell Signaling Technology; Cat. no. 23064; dilution 1:400). After the incubation, cells were washed with PBS three times, each time for 5 minutes. Secondary antibodies Alexa Fluor 488 donkey anti-mouse (Thermo Fisher Scientific; Cat. no. A-21202) and Alexa Fluor 555 donkey anti-rabbit (Thermo Fisher Scientific; Cat. no. A-31572) were diluted (1:500) in 1% BSA in PBS and applied for 2 hours at RT in the dark. Cells were then washed with PBS three times, each time for 5 minutes, incubated in 300 ng/ml DAPI in PBS for 10 minutes, and rinsed three times with PBS, each time for 5 minutes. Finally, samples were later visualized using a fluorescent microscope (Olympus; Cat. no. IX73).

### Statistical Analyses

Fisher’s exact test was used for comparison of amplification frequency and incidence of large deletions in PCR amplicons shown in **Fig. 2c**, f. Fisher’s exact test was used for comparison of frequencies of large deletions at *PCSK9* and *HBG1/2* target loci in NGS reads from pooled CRISPR- and ABE-treated samples shown in **Extended Data Fig. 2a, b**. A non-parametric *t*-test was used for comparison of gene densities around chromosome breaks in ABE-treated embryos and random genomic loci shown in **Fig. 3f**. One-way ANOVA was used for comparison of developmental potential of human embryos injected with ABE mRNA or ABE RNP at the 2PN stage or at fertilization, shown in **Fig. 4b**. A *p*-value less than 0.05 was considered significant. GraphPad Prism (Version 10.3.1) was used to calculate all plotted statistical comparisons (* *p* ≤ 0.05, ** *p* ≤ 0.01, *** *p* ≤ 0.001, and **** *p* ≤ 0.0001). No statistical methods were applied to predetermine sample sizes, as human embryo samples are difficult to project due to limited availability.

## ACKNOWLEDGMENTS

This work was supported by the Institute of Organic Chemistry and Biochemistry (IOCB)–Tech Foundation research grant, the New York Stem Cell Foundation (NYSCF)–Druckenmiller Advanced Postdoctoral Fellowship, and the resources of Genomic Prediction, Inc. in chromosomal analysis. Research in the Bae lab was supported by the Korean Fund for Regenerative Medicine (KFRM) No. RS-2024-00332601 to S. Bae.

## AUTHOR CONTRIBUTIONS

S.J. designed the study and performed editing analysis and chromosome analysis. J.K. contributed to editing in cultured cells, genotyping, and editing analysis. J.S. performed mapping of chromosome breaks. C.J. conducted the Digenome-seq experiments and analysis. M.I.R.K. contributed to mRNA synthesis, mapping of chromosome breaks, and embryology. H.J. and J.W. provided purified base-editing proteins from human cells. M.I. performed large deletion analysis and contributed to genotyping. M.L. performed a Western blot. S.Bh. performed immunostaining. M.K. contributed to mRNA synthesis and analysis of embryo development. S.X. performed gene density analysis at break sites. G.H. assisted with Digenome-seq data analysis. S.Ba. contributed to the design of off-target analysis. D.M., J.X., and N.T. performed WGA and SNP array analysis. D.E. performed embryology and contributed to all aspects of the study, including design, data analysis, and manuscript writing. S.J. wrote the manuscript, and all authors contributed to the editing of the manuscript.

## DATA AVAILABILITY

SNP array data are available at Gene Expression Omnibus (GEO) under the accession no. GSE290961 (https://www.ncbi.nlm.nih.gov/geo/query/acc.cgi?acc=GSE290961). The following secure token has been created to allow review of record GSE290961 while it remains in private status: **utcdsukkfvwbtkn**. Donors of gametes have provided informed consent for genetic analysis. Data sharing was approved by the Institutional Review Board based on de-identification and the type of genetic information content.

## CODE AVAILABILITY

This study did not generate a code. Genomic Prediction, Inc. provides preimplantation embryo sample analysis service using gSUITE software.

## MATERIALS & CORRESPONDENCE

hESC lines derived in this study are available upon request from the Lead Contact.

Please direct requests for further information, reagents, and resources to the Lead Contact, Dieter M. Egli

